# Cultural variation in voting patterns reflects the landscape of US genetic diversity: A test of the cultural niche hypothesis

**DOI:** 10.1101/856450

**Authors:** Jason A. Hodgson

## Abstract

Humans are unique in the animal world in the extent to which we have the potential to affect our own biological evolution through culturally mediated behavioural variation. However, there is scant clear evidence of the phenomenon. Here, I estimate genetic diversity for every county in the US by merging demographic data from the 2010 US Census with genomic reference populations and show that genetic diversity is predicted by cultural variation as reflected by the 2016 Presidential Election. Remarkably, the 2016 Election alone explains 18% of US county-level variation in genetic diversity, with diversity decreasing as Trump support increases. When state level variation is included in the model, 71% of variation is explained. Within states, genetic segregation increases with cultural segregation. I then tested whether the observed genetic patterns might be explained by culture acting on race and ethnicity and found that genetic diversity tends to increase with racial and ethnic diversity, and that the 2016 Election is an even better predictor of the US landscape of ethnic diversity. Finally, I measured patterns of ethnic assortative mating nationwide from the Census data, and found that ethnic assortment is primarily explained by the proportion of minorities in each county, and that the most diverse counties are the most assortative. Overall, regional variation in cultural tolerance appears to be structuring biological diversity on a massive scale. Also, assortative mating is maintaining ethnic and genetic diversity within the most diverse and tolerant areas. Culture is a primary driver of biological evolution across the US.

## INTRODUCTION

Two of the major outstanding questions in the study of human evolution are 1) what is the extent and trajectory of our on going evolution, and 2) to what extent is culture influencing the course of our evolution? Both of these questions have been largely neglected or even actively avoided by the research community in favour of a focus on our past history of evolution. Indeed, most genetics studies of human evolution using employ a sampling strategy designed to capture the diversity present several generations ago, no matter the diversity that exists today. For example, to investigate the population history of Great Britain, the British Isles Project sampled people from across the island for which all four grandparents were born within 60km of each other in the local area [1, 2]. This strategy likely captured British diversity from several generations, however because the sampling criteria describe very few people today it will not capture current diversity. Similar sampling strategies have been employed in almost all surveys of human genetic diversity. Researchers typically include only people who have all four grandparents of the researcher’s desired ethnic group no matter the actual ethnic and demographic mix of region ([e.g. 3, 4]). For this reason, and though it is seldom acknowledged, very little is known about the actual current distribution of human genetic diversity ([for an exception see 5]). In most parts of the world, the current distribution of human genetic diversity is likely to reflect the layered and complex histories of migrations into the area, as well as the extent to which ethnic divisions are preserved through assortative mating [6, 7]. Human cultures vary in how open they are to outsiders, and also in rates of acculturation (the process of adopting the practices and beliefs of the local population after migration), and these parameters are known to affect both migration itself and the rate of cultural homogenisation [8]. If there is genetic structure between ethnic or cultural groups, these cultural phenomena are expected to also variably affect regional genetic diversity. It may very well be the case that cultural variation is a primary driver of the current distribution of genetic diversity.

Gene-culture coevolution is the idea that both allele frequencies and culturally mediated behavioural variation can influence change in each other and therefore evolve in concert [9–11]. Classic examples include cultures that practice cattle herding being associated with increased frequency of the allele associated with the ability to digest milk sugar throughout adulthood (lactase persistence) [12], and cultures that practice cereal agriculture being associated with greater copy numbers of a gene that promotes the ability to digest dietary starch [13]. It has also recently been shown that the Bajou people of Southeast Asia who practice a unique form of fishing through breath hold diving also have increased frequencies of genes associated with large spleens that act as a reservoir of oxygenated red blood cells [14]. These examples involve a feedback mechanism where the cultural behaviour selects for the increase in a specific gene variant that in turn better enables the cultural behaviour [10], and the effects are therefore localised to specific sites in the genome. It is also possible for culturally mediated behavioural variation to affect allele frequencies across the entirety of the genome if the cultural variation promotes or inhibits the association of people from a wider or narrower range of human diversity. The amount of genomic diversity across the human species range has been explained geographically, with genetic diversity decreasing with increasing distance from East Africa [15–17]. This loss of genetic diversity is also associated with a corresponding loss of phenotypic diversity [18–20]. The reduction in genetic and phenotypic diversity is thought to be a consequence of the reduction in population size due to the serial founder effect as anatomically modern peoples spread out of Africa in the late Pleistocene [17, 18]. These studies describe how ancient evolutionary forces affected the landscape of genetic and phenotypic diversity that existed prior to the modern era. However, mass migration associated with e.g. colonialism, slavery, and differential population expansion has largely erased the association between geography and diversity that once existed. It is possible that cultural variation is now a primary driver of regional variation in genetic diversity. There is tremendous variation between cultures in the extent to which they are open or closed to diversity of all kinds [21]. The cultural niche hypothesis predicts that cultural endogamy should decrease genetic diversity and exogamy should increase it.

The 2016 US Presidential Election offers the opportunity to test the cultural niche hypothesis on a massive scale. US election results are known to reflect regional cultural variation, though the cultural variables being reflected are often particular to each election [22, 23]. There is evidence for regional variation in tolerance for diversity within the US [24]. Because Donald Trump largely ran on a platform of cultural and national insularity, with key campaign promises being to wall off the US southern border, and to ban Muslims from entering the country [25], it is expected that the 2016 US Election results capture regional variation in tolerance for diversity. Indeed, support for Trump is thought to coincide with pervasive racism [26]. Thus, though there is undoubtedly variation among individuals in their reasons for supporting a particular political candidate, regional variation in support for Donald Trump should capture regional variation in cultural tolerance of diversity to a large degree. The cultural niche hypothesis then predicts that greater regional support for Trump will correspond to regions with reduced genetic diversity. Exit polling following the 2016 election found that racial and ethnic minorities overwhelmingly preferred Hillary Clinton [27], and this is supported by post-election analysis which shows race to be a primary predictor of regional election results[28]. If ethnic diversity is associated with increased genetic diversity, and is geographically structured, the cultural niche hypothesis is likely to be supported. I test this with a study design that incorporates almost the entire current population of the United States (322,649,933 people).

## RESULTS AND DISCUSSION

The US Census is taken every 10 years and records demographic data including racial and ethnic affiliation for every individual in the US regardless of legal status. These data make it possible to estimate genetic diversity for each US county, by calculating county level allele frequencies as the mean allele frequency of genetic reference samples for each reported US Census ethnic category, weighted by the demographic proportion of each ethnic group in the county. I used data available from the 1000 Genomes and Human Genome Diversity Projects [3, 4, 29] to represent the allele frequency variation for each of the US Census ethnic categories of Hispanic or non-Hispanic ‘white’, ‘African American’, ‘Asian’, ‘Native American’, or ‘Pacific Islander’. Allele frequencies were estimated for each county for an unbiased [29] and unlinked panel of 46,155 single nucleotide polymorphisms (SNPs) from which genetic diversity (*H_e_*) was calculated as the mean expected heterozygosity [30] (see methods).

I used election data from the MIT Election Data + Science lab [31], which provides county-level results from the 2016 Presidential election for 3,113 counties (or their equivalents). This represents the entirety of the US with the exception of Alaska (for which county level reporting is not available). I compared these data to the estimates of county level genetic diversity. The data are summarised in **Table 1.** The distribution of county-level voting is skewed towards Trump (***Supporting information*, Figure S1**). Trump won 84% of counties (2,623 to 488) but remarkably 55% of the US population lives in counties won by Clinton – an excess of 30,695,723 people – highlighting the tenfold advantage in population densities in Clinton counties. The genetic reference populations have genetic diversities as follows: European American (*H_e_* = 0.215), African American (*H_e_* = 0.269), Hispanic American (*H_e_* = 0.219), Native American (*H_e_* = 0.181), Asian American (*H_e_* = 0.221), and ‘Pacific Islander’ (*H_e_* = 0.202). The county level average genetic diversity is 0.225 and ranges from 0.188 to 0.267, with Oglala Dakota County, South Dakota being the least diverse, and Clairborne County, Mississippi being the most diverse. The distribution of county-level genetic diversities is highly skewed with most counties clustered just above the genetic diversity of the European American reference population and a heavy right tail extending towards the diversity of the African American reference (***Supporting information*, Figure S2**). There are also a small minority of counties in the left tail in the direction of the diversity of the Native American reference.

**Table 1.** Summary of data included in this study. Genetic diversity (*H_e_*) is expressed as the unweighted mean and standard deviation of each county, as well as the means and standard deviations weighted by county population size.

|  | Clinton | Trump | USA |
| --- | --- | --- | --- |
| **Counties** | 488 (16%) | 2,623 (84%) | 3,113 |
| **Votes** | 65,623,888 (51%) | 62,790,048 (49%) | 128,413,936 |
| **Population mean** | 362,034 | 55,653 | 103,713 |
| **Population S.D.** | 730,673 | 131,777 | 332,657 |
| **Total population** | 176,672,828 (55%) | 145,977,105 (45%) | 322,649,933 |
| **Area mean (sq. mi.)** | 1,133 | 982 | 1,005 |
| **Area S.D. (sq. mi.)** | 1,761 | 1,232 | 1,330 |
| **Total Area (sq. mi.)** | 553,434 (15%) | 3,211,900 (85%) | 3,765,387 |
| **People per sq. mile mean** | 972 | 90 | 228 |
| **People per sq. mile S.D.** | 3,163 | 189 | 1,302 |
| **Genetic diversity ($H_e$)** | 0.234 | 0.222 | 0.225 |
| **$H_e$ S.D.** | 0.016 | 0.008 | 0.011 |
| **Weighted $H_e$** | 0.233 | 0.225 | 0.230 |
| **Weighted $H_e$ S.D.** | 0.010 | 0.008 | 0.010 |

### US genetic diversity increases with ethnic diversity

I next explored the relationship between demography and genetic diversity. There is clear geographic structuring of ethnicity in the US, with African Americans clustered in the East and throughout the Old South, Hispanic Americans clustered in the Southwest, Florida, and Eastern cities, Asian Americans largely dotted throughout the major population centres, Native Americans in pockets of the West and Southwest, and Pacific Islanders largely restricted to Hawaii (**Figure 1**). The relationship between county demography and genetic diversity was investigated by plotting the empirical expectation of genetic diversity for two-way demographic mixes of each of the minority US Census ethnic groups and the majority European Americans (see methods). Genetic diversity is maximised at intermediate ratios of the minority group and European Americans for each ethnic group except African Americans, for which genetic diversity continues to increase until the population is entirely African American (**Figure 2**). Each US county was plotted according to their respective proportions of minority ethnicities relative to European Americans and their genetic diversities. For most counties greater genetic diversity is associated with demographics incorporating a greater proportion of minority ethnic groups. The counties with the highest genetic diversity are predominantly African American, while the counties with the lowest genetic diversity are predominantly Native American.

**Figure 1.**
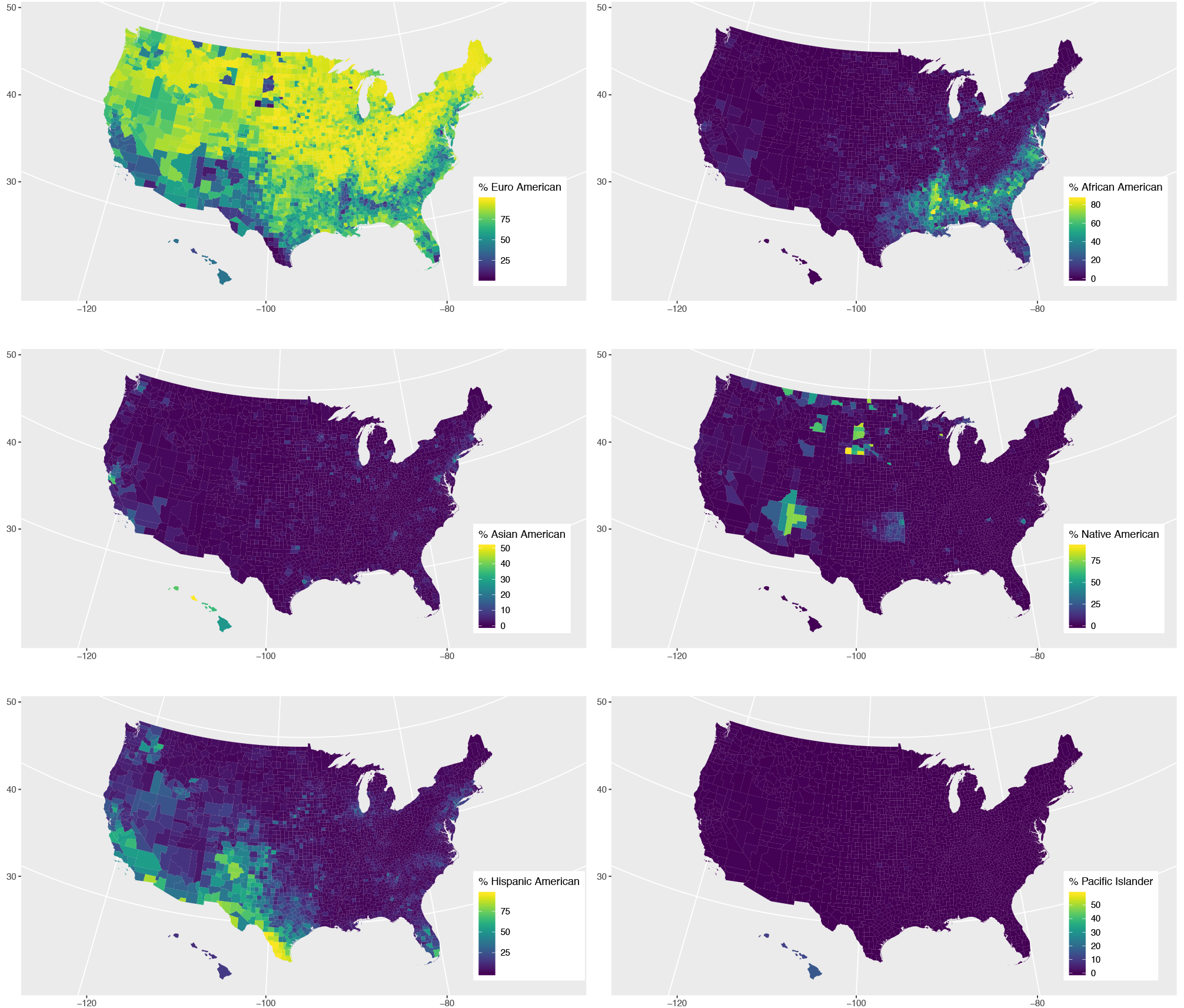
The US ethnic landscape. County level proportions of each ethnic group included in the 2010 US Census for the year 2016.

**Figure 2.**
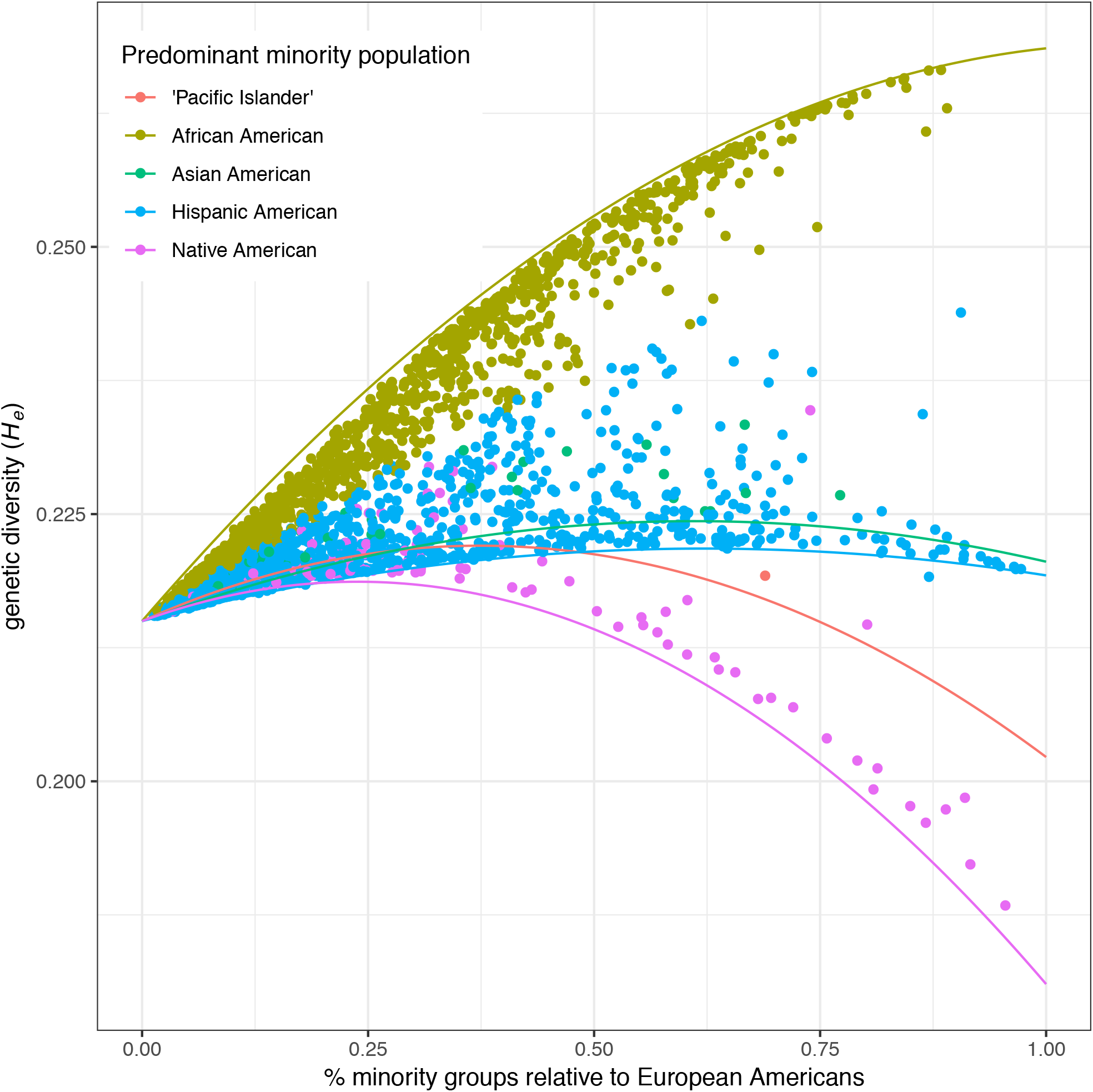
The relationship between US genetic diversity and demography. The curves indicate the theoretical genetic diversities for a population consisting of a two-way mix of European Americans and the indicated minority group at various demographic ratios (see methods). Genetic diversity is maximised at intermediate demographic ratios for all minority groups except African Americans for who genetic diversity continues to increase as their proportion in the population increases. The points represent each US county coloured by the predominant minority population in the county. Counties with the greatest proportion of African Americans have the greatest genetic diversity.

### As support for Donald Trump increases genetic diversity decreases

As expected from the cultural niche hypothesis, counties that voted for Trump have lower average genetic diversity than counties that voted for Clinton (0.222 to 0.234) (**Table 1**). A plot of county-level Trump vote percentage versus genetic diversity shows that as Trump voting increases genetic diversity decreases (***Supporting information*, Figure S3**). Linear regression found that Trump vote percentage explains 18% of county-level genetic diversity nationwide (F_1,3109_=689.1, adj. R^2^=0.181, p<2.2e-16, AIC=8751.5), and that a 10% increase in Trump voting is associated with a 0.3% decrease in genetic diversity. The magnitude of this relationship is remarkable given the small range of variation of genetic diversity nationwide (18.8% to 26.7%). These findings indeed suggest that as cultural tolerance increases so does genetic diversity.

The landscapes of cultural and genetic diversity were further explored by plotting the maps of each. Counties of high genetic diversity are clustered primarily in the Cotton Belt of the Old South [32] (**Figure 3a**) in counties with high proportions of African Americans (**Figure 2**), reflecting America’s legacy of slavery. Secondary pockets of high genetic diversity correspond to America’s older Eastern cities. Areas with the lowest genetic diversity are found in Western and Southwestern counties where Native Americans are in high proportion. These findings broadly agree with the landscape of genetic diversity inferred from 23andMe customers [5]. The landscape of cultural tolerance of diversity as reflected in Trump voting percentage largely mirrors the landscape of genetic diversity with blue (Clinton) areas of the map corresponding to the high genetic diversity areas of the Old South and the Eastern cities, however there are some apparent discrepancies in counties in Western states that show preference for Clinton but not particularly high genetic diversity (**Figure 3b**). There is known to be state-level cultural variation in the US [33], and there is also state-level variation in ethnic diversity. It is possible that cultural variation in tolerance of diversity affects regional patterns of genetic diversity according to the state-level diversity environment. To test for this I created a linear model to predict genetic diversity from voting patterns incorporating state-level variation. This model explains 71% of variation in genetic diversity nationwide (F_50,3060_=151.6, adj. R^2^=0.708, p<2.2e-16, AIC=5595.2), and has far greater predictive power than a model that considers state environment alone (F_49, 3063_=79.8, adj. R^2^=0.554, p<2.2e-16, AIC=6925.3). The predicted landscape of genetic diversity from this model shows remarkable concordance with actual genetic diversity, with the primary exceptions being the counties with majority Native American ethnicities (**Figure 3c,d**).

**Figure 3.**
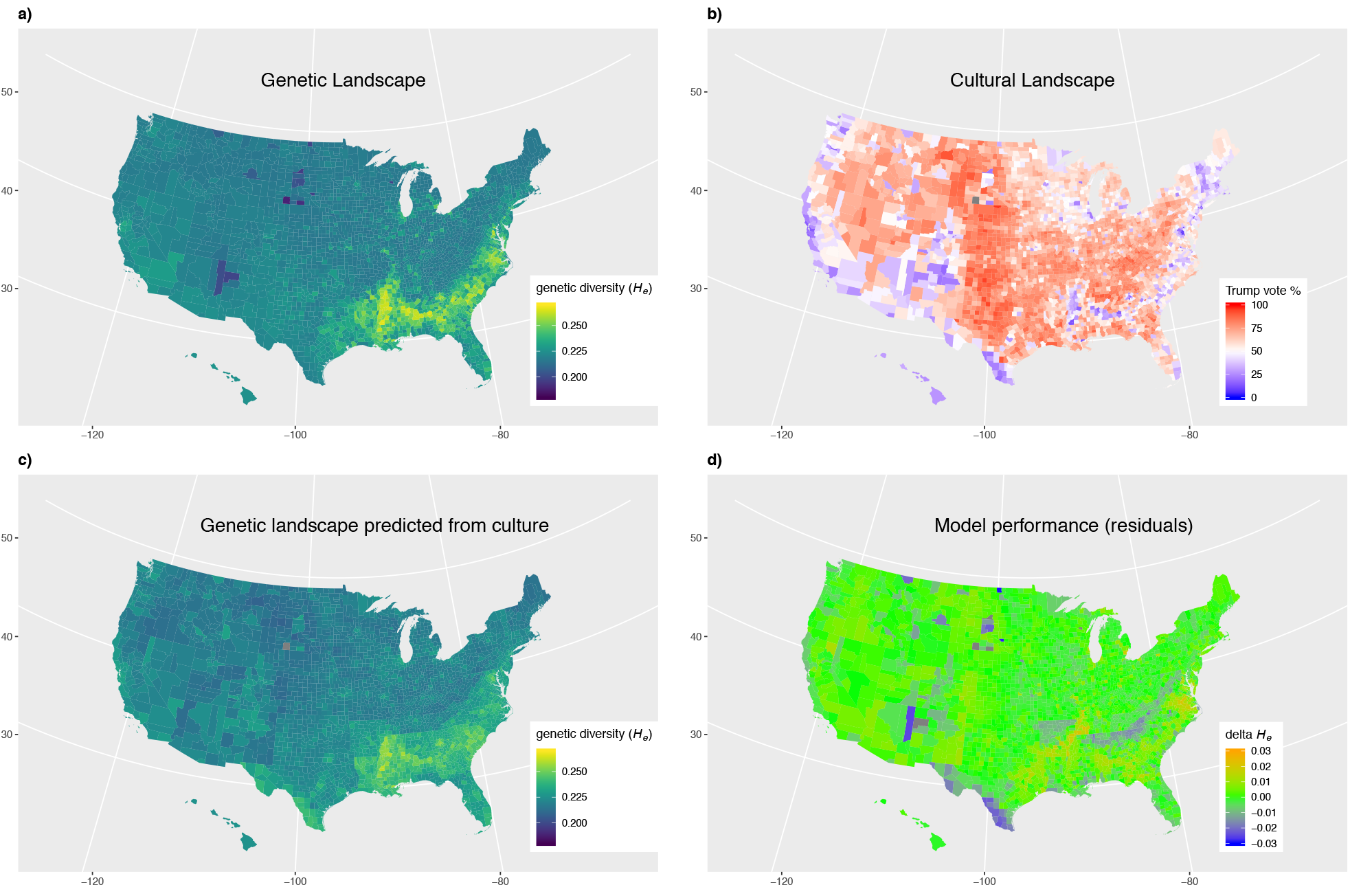
Landscapes of genetic and cultural diversity. **a)** Genetic diversity (*He*) for each county. **b)** Cultural tolerance of diversity as indicated by 2016 election results for each county. **c)** Genetic diversity for each county predicted from a linear model that considered the percent of the vote for Donald Trump, and the State of each county (F=151.6_50, 3060_, adj. R^2^=0.708, p<2.2e-16, AIC=5595.2). **d)** The difference between observed and model predicted genetic diversity for each county. Green indicates the model is accurately predicting genetic diversity, blue indicates the model is overestimating, and orange indicates the model is underestimating genetic diversity.

### The most culturally divided states are also the most genetically divided

These findings indicate that counties that voted for Trump tend to have lower genetic diversity than counties that voted for Clinton no matter the overall level of diversity of the state. To further explore this I plotted the mean genetic diversities for counties won by either Trump or Clinton for each state (**Figure 4a**), as well as their density distributions (***Supporting information*, Figure S4**). Thirty-nine states have greater genetic diversity in counties Clinton won, compared to only seven states with greater diversity in Trump counties. The remaining four states were won entirely by either Clinton or Trump. Nationwide, the vast majority of Trump counties have genetic diversity similar to the European American reference. The difference in diversity between Trump and Clinton counties tends to increases as the overall diversity of the state increases. If culture is driving this observed genetic differentiation then it is expected that the most culturally divided states should also be the most genetically divided. To test this, I calculated indices of cultural (*Q_C_*) and genetic (*Q_G_*) niche separation for each state (see methods). These indices express the niche separation for each state in units of standard deviations of the total variation within the state, and are thus standardised to the overall environment of the state. As predicted, genetic niche separation increases with greater cultural niche separation (**Figure 4b**), and cultural niche separation explains 47% of the variation in genetic niche separation (F_1,43_=40.7, adj. R^2^=0.47, p=1.03e-07). The most genetically and culturally segregated states are Missouri, Kansas, Tennessee, and Kentucky. Idaho, Wyoming, and Texas are notable for being highly culturally divided but not genetically divided. Idaho and Wyoming are both ethnically homogenous with predominant European American populations, while Texas is ethnically diverse. Trump only lost Latah and Blaine out of Idaho’s 44 counties. Latah County is home to the University of Idaho, while Blaine is the home of a major ski resort and consequently median home prices that are more than double the state average. Similarly, Trump won all of Wyoming’s 23 counties except Teton County, which is far more expensive than the rest of the state and contains the exclusive Jackson Hole ski area. American populations with university level education have been found to be more culturally tolerant [24], and education may explain the cultural division between the wealthy areas and university towns in these states. Trump lost 27 of Texas’s 254 counties, all of which are ethnically diverse but with the predominant minority group being Hispanic Americans. The lack of separation in genetic diversity despite the cultural division is explained by Hispanic Americans having similar genetic diversity to European Americans.

**Figure 4.**
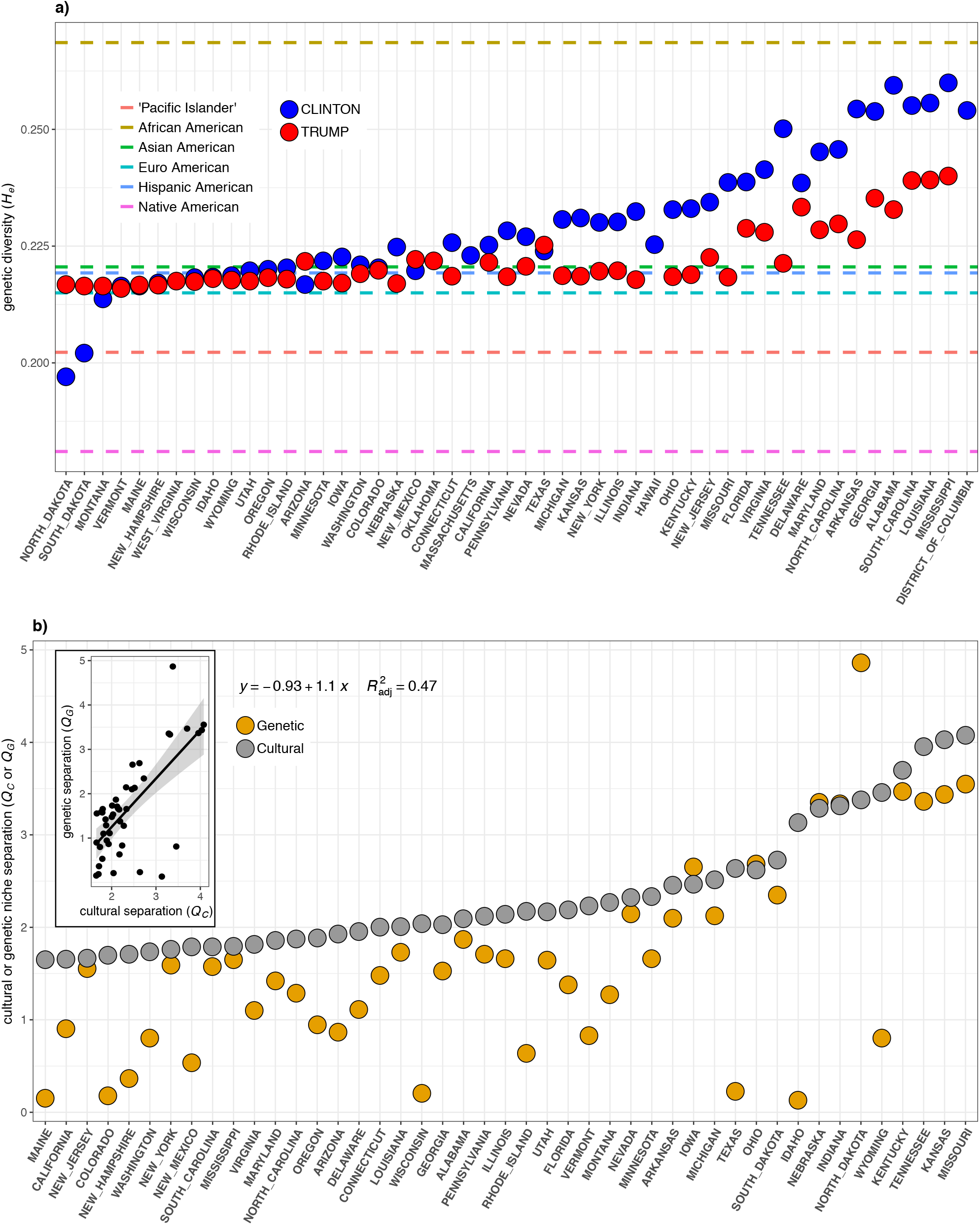
Genetic and cultural separation within states. **a)** The relationship between mean genetic diversity (*H_e_*) for counties won by either Donald Trump or Hillary Clinton for every state in the US except Alaska (for which county-level voting results are not available). The dashed lines indicate the genetic diversities for the genomic reference populations used to estimate county-level diversities according to their particular demographies. For most states Trump counties have genetic diversity slightly above that of European Americans consistent with their homogenous-demographics. Thirty-nine states have greater diversity in Clinton counties, five states were entirely won by a single candidate, and only seven states have higher diversity in Trump counties. These are primarily Western states where the majority of the ethnic diversity is Native American. **b)** The relationship between cultural (*Q_C_*) and genetic (*Q_G_*) niche separation in all states that were not won entirely by one candidate. For most states, cultural separation is greater than genetic separation. The inset shows the same data as a scatter plot. There is a strong positive relationship between cultural and genetic niche separation (F_1,43_=40.7, adj. R^2^=0.47, p=1.03e-07).

### The 2016 Election is a better predictor of ethnic diversity than genetic diversity

As predicted, US genetic diversity is lower in culturally less tolerant areas, suggesting that cultural variation can impact biological diversity on a massive scale. However, because genetic diversity is not readily apparent or recognised by people it is unlikely that the mechanism driving the pattern is preference related to biological diversity per se. The Trump campaign’s rhetoric targeted ethnic minorities including Hispanic and Muslim populations [25], and since then Trump has continued to promote division between the majority European Americans and minority groups by invoking classic racist tropes when suggesting that African American and Muslim American US congresswomen go back to the countries from which they came (despite them being American-born) [34]. This suggests that the mechanism driving the differentiation of genetic diversity may be relative lack of tolerance of ethnic diversity in communities that are predominantly European American. Because genetic diversity tends to increase as the proportion of minority groups increase (**Figure 1**), if communities are structured according to tolerance for ethnic diversity, a by product would be low genetic diversity in predominantly European American communities and greater diversity in more ethnically diverse communities. If so, it is expected that cultural variation will be a better predictor of ethnic diversity than it is of genetic diversity. To test this, I plotted the percentage Trump vote against the percentage of European Americans in each county and found that as Trump voting increases counties tend to get more predominantly European American (***Supporting information*, Figure S5**). Linear regression found that Trump voting explains 28% of the variation, and every 10% increase in Trump voting is associated with a 6.7% increase in the percentage of European Americans (F_1,3109_=1192, R^2^=0.28, p<2.2e-16, AIC=26,468.1). When state level variation is included in the model, Trump voting explains 73% of the variation in percentage of European Americans (F_50,3060_=170.8, R^2^=0.73, p<2.2e-16, AIC=23,429.9), and this model does a remarkable job predicting the landscape of ethnic diversity in the US (**Figure 5**). Thus, culture does explain slightly more variation in ethnic diversity than genetic diversity suggesting that opposition to ethnic diversity (i.e. racism) may be the causal mechanism driving the pronounced differentiation of genetic diversity across the US.

**Figure 5.**
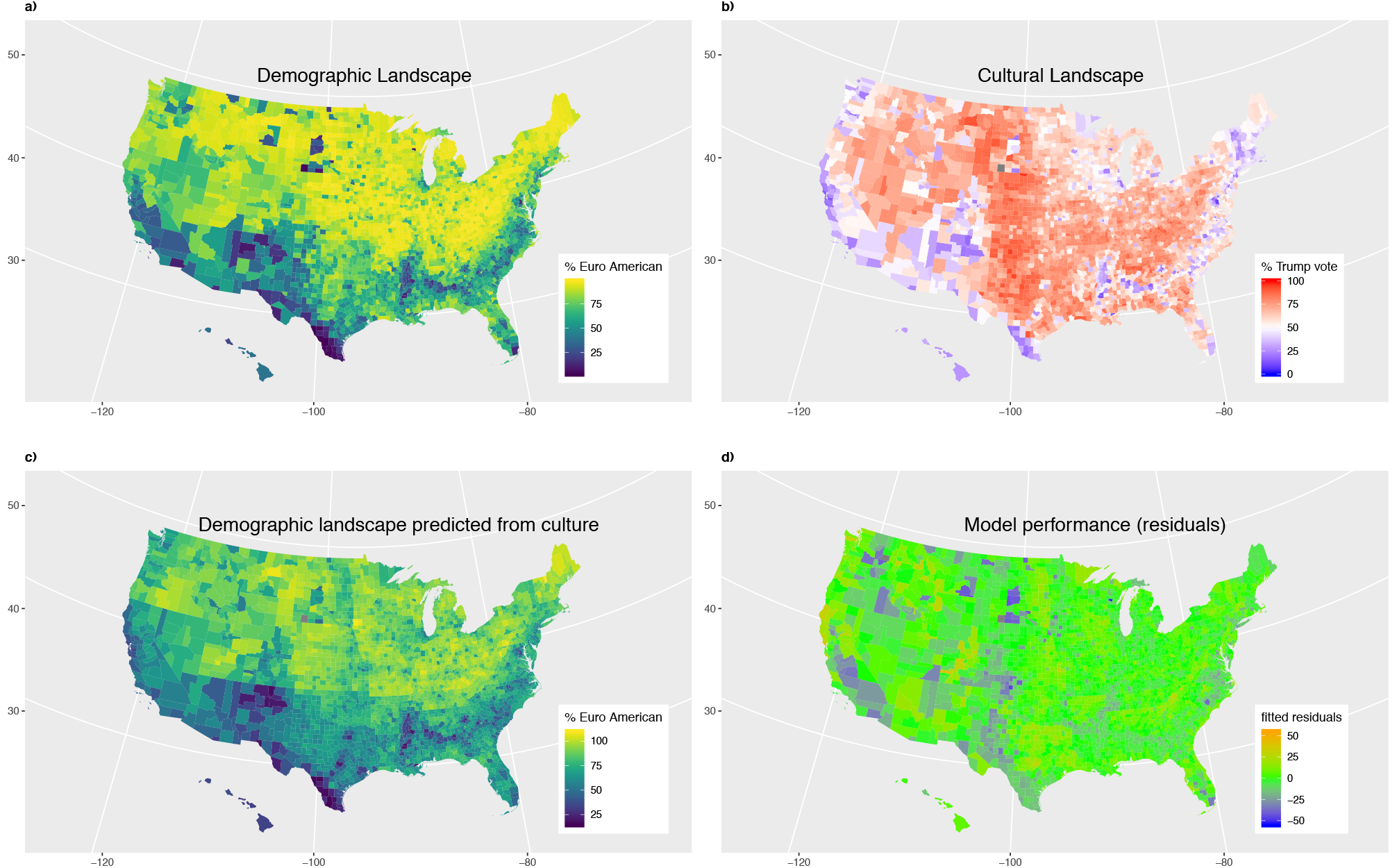
Landscapes of demographic and cultural diversity. **a)** The demographic landscape as indicated by the percentage of the majority ethnic group - European Americans - for each county. **b)** The landscape of cultural tolerance of diversity as indicated by 2016 election results for each county. **c)** The percentage of European Americans for each county predicted from a linear model that considered the percent of the vote for Donald Trump, and the US State for each county (F=170.8_50, 3060_, adj. R^2^=0.73, p<2.2e-16, AIC=23,429.9). **d)** The difference between observed and model predicted percentage of European Americans for each county. Green indicates the model is accurately predicting the demographics, blue indicates the model is overestimating, and orange indicates the model is underestimating the percentage of European Americans.

### Population density does not explain the landscape of US genetic diversity

Because of the tenfold greater population density in counties won by Clinton, it is possible that the landscape of genetic diversity is actually being driven by population density rather than cultural variation per se. To test this, I used linear regression to predict genetic diversity from log10 population density and found a positive relationship (F_1,3109_=292.2, R^2^=0.086, p<2.2e-16, AIC=9095.0) (***Supporting information*, Figure S6**). However, population density explains less of the variation in genetic diversity than does Trump voting percentage (9% versus 18%). I then fit a model that predicts genetic diversity from population density after controlling for state level variation and found that this model performs only marginally better than the model that uses state level variation alone (F_50,3060_=84.9, R^2^=0.574, p<2.2e-16, AIC=6766.1) (***Supporting information*, Figure S7**). This is likely because state level variation in population density captures most of the explanatory power of population density variation itself. The five models used to predict US genetic diversity in this study were compared using the Akaike Information Criterion (AIC) (**Table 2**), and the model that used cultural environment as measured by Trump vote percentage controlled for state was substantially better than each of the other models [35]. These findings suggest that variation in cultural environment, irrespective of population density is indeed influencing the landscape of US genetic diversity.

**Table 2.**
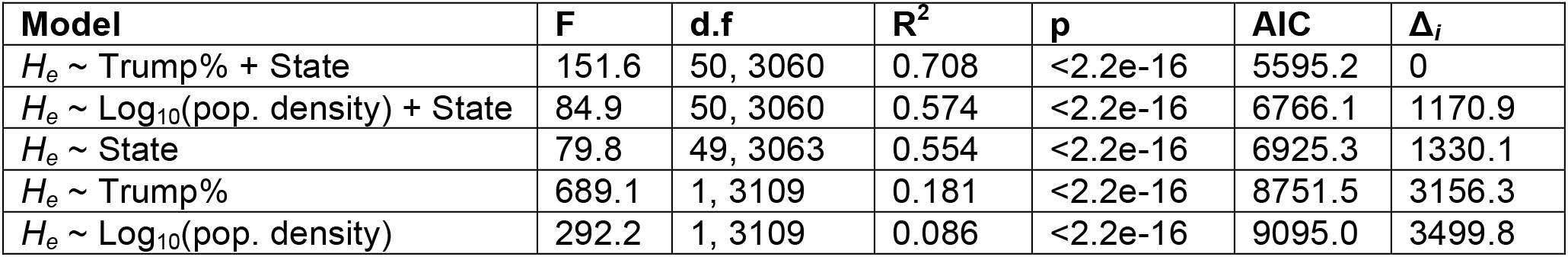
Comparison of linear models used to predict US county-level genetic diversity (*H_e_*). The Akaike Information Criterion (AIC) was used to compare model performance. The model that best explains genetic diversity considers Trump vote percentage and state. The Δ_*i*_ shows the increase in AIC relative to this best model.

### Patterns of assortative mating suggest ethnic divisions will persist even in diverse and tolerant areas

The findings presented above strongly suggest that US culture is structuring the landscape of ethnic diversity, which in turn is structuring the landscape of genetic diversity. Because ethnic diversity is geographically structured into communities there is the potential for this diversity to break down through time through intermarriage [8]. The extent to which people tend to mate within or between ethnic groups will thus determine the extent to which current ethnic divisions will persist into the future. The US Census provides information on the number of people in each US county with mixed ethnicity. I used this information to calculate an index of assortative mating (*A*) that describes the extent to which mating patterns diverged from randomness with respect to ethnicity. The index of assortative mating is expressed as the proportion of mate pairs that are assortative (positive values) or disassortative (negative values). Thus, values of *A* equal to one indicate 100% assortative mating, values equal to negative one indicate 100% disassortative mating, and values near zero indicate random mating. Unexpectedly, there is a weak negative relationship between Trump vote percent and *A*, with mating patterns in areas that support Trump being less assortative (F_1,3109_=198.8, R^2^=0.06, p<2.2e-16) (***Supporting information*, Figure S8**). This pattern could be explained if high levels of ethnic assortative mating require a sufficiently large minority population for there to be a sufficient number of available mates within groups. To test this, I plotted assortative mating against the proportion of minorities for each US County (**Figure 6**). There is a clear nonlinear relationship, with assortative mating rapidly increasing as the minority population becomes less rare, and plateauing at near perfect assortative mating when the minorities reach about 25% of the population. A fourth order polynomial regression found that the proportion of minorities explains 55% of variation in assortative mating index nationwide (F_4,3108_=935.6, R^2^=0.55, p<2.2e-16). Counties where the predominant minority groups are Native American or Asian American show relatively less assortative mating. Hawaii County, Hawaii shows remarkably high ethnic disassortative mating with a mating index of *A* = −0.43. In this county it is expected that 21% of people would have mixed ethnicity if mating were random, however 30% of the 198,681 people report a mixed background. Overall, ethnic assortative mating predominates throughout the US and this pattern largely depends on the local size of ethnic minority communities, rather than the cultural environment. Assortative mating is thus maintaining ethnic diversity in the most diverse regions of the US. Assortative mating is known theoretically to also maintain genetic and phenotypic diversity [36, 37]. If the observed patterns of assortative mating persist into the future, the current ethnic and genetic landscapes are expected to persist as well.

**Figure 6.**
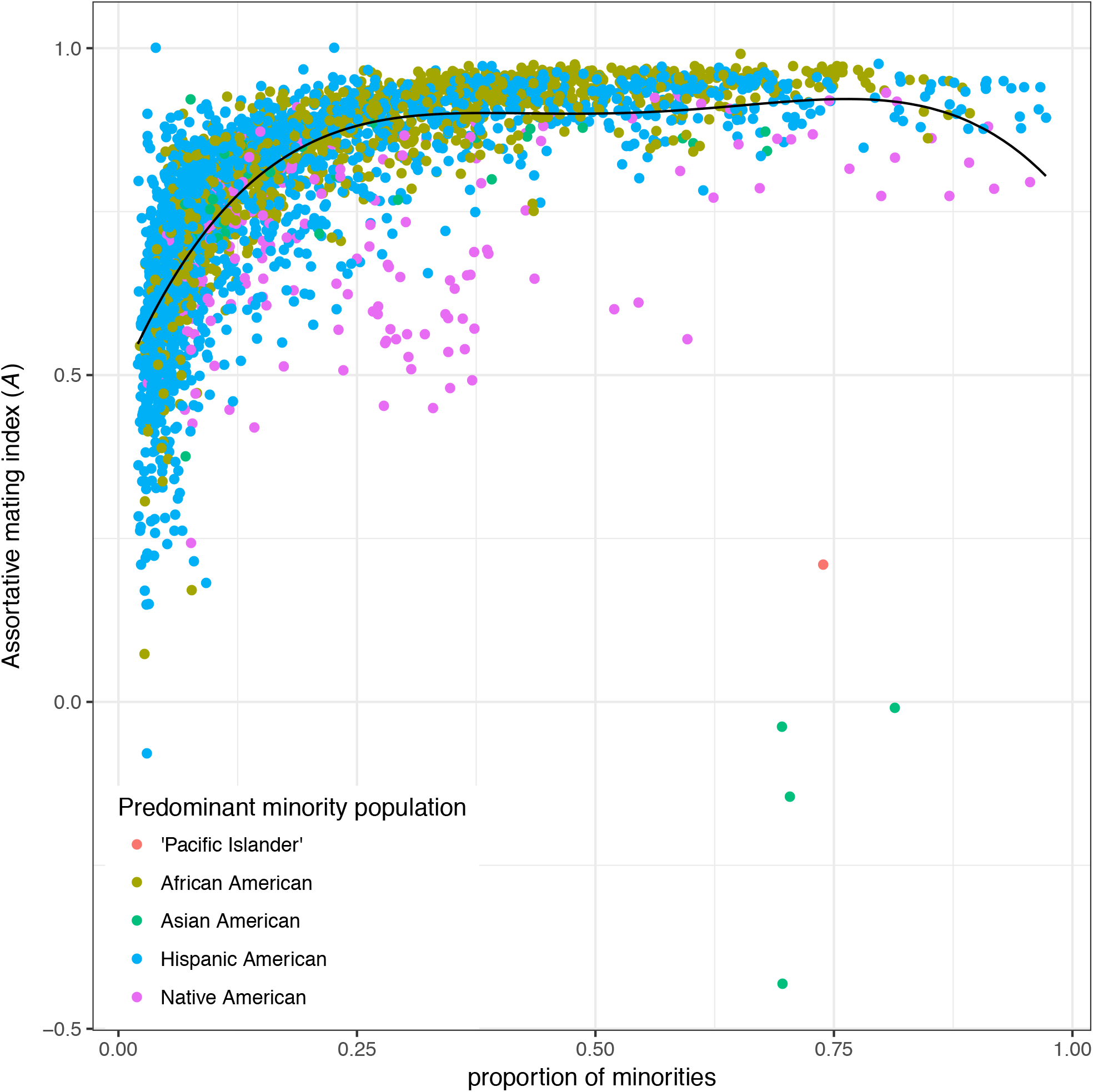
The relationship between assortative mating and ethnic demography. The assortative mating index, *A*, for each US county is plotted against the proportion of minorities. Each county is coloured according to its predominant minority group. The best-fit curve from a fourth order polynomial regression is shown. Proportion of minorities explains 55% of assortative mating nationwide (F_4,3108_=935.6, R^2^=0.55, p<2.2e-16), and assortative mating is maximised once minorities reach ~25% of the population.

### Broader implications

This research has found that 1) genetic diversity tends to increase with ethnic diversity, 2) the cultural landscape of tolerance of diversity predicts the landscapes of ethnic and genetic diversity, 3) the most culturally divided states also tend to be the most genetically divided, and 4) assortative mating is maintaining ethnic diversity even in the most culturally tolerant areas. These findings strongly support the cultural niche hypothesis that variation in cultural tolerance for diversity is influencing the landscape of genetic diversity across the US. Racial and ethnic groups in America are socially – rather than biologically – defined [38]. However, there are allele frequency differences between these social groups [39], and social forces that work on race or ethnicity will have biological effects. Ethnic minorities in the US tend to live in areas that are more culturally tolerant of diversity and this in turn increases the genetic diversity of culturally tolerant areas, and decreases it in culturally intolerant areas. The most genetically diverse areas of the US were found to correspond to the Cotton Belt of the Old South [33] and is explained by high proportions of African Americans in these counties. These counties reflect the legacy of race-based chattel slavery that ended in 1865 and was followed by legally enforced segregation during the Jim Crow era until the Civil Rights act of 1964 [40]. Since then, racism has been a persistent problem in America [26]. This suggests that culture has long been shaping the landscape of US genetic diversity, and that the patterns observed today have deep roots within the American cultural milieu. Remarkably, European Americans who currently live in counties that had high concentrations of slaves in 1860 have been found to be more racist than European Americans from elsewhere in the South [41]. These attitudes might affect migration patterns in two ways that would reinforce the patterns found in this study. First, African Americans in these areas may be dissuaded from moving to areas with a higher proportion of European Americans because their local experience of them is negative. Second, European Americans with these attitudes might migrate towards less diverse areas, thus increasing the cultural and ethnic segregation. Future research should directly test the causative nature of the relationship between cultural and genetic diversity. It is predicted that patterns of migration within the US will track cultural tolerance, with ethnic minorities tending to move to culturally tolerant areas, and culturally intolerant European Americans moving to more homogenous areas.

Surprisingly, assortative mating with respect to ethnicity was not found to be strongly related to cultural tolerance. Instead, the primary predictor of ethnic assortative mating is the proportion of minorities in a county, with assortative mating typically exceeding 90% of mate pairs once minorities reach 25% of the county. This finding has major implications for the maintenance of racial and ethnic divisions within the US [8], as well as for the maintenance of genetic diversity [36, 37]. As expected, the majority of minorities in the US reside within more culturally tolerant areas, however this cultural tolerance does not appear to extend to mate choice patterns. Race and ethnicity have historically functioned like caste divisions in the US, and these group divisions can still be observed in social interactions [42]. There are extensive inequalities between US racial and ethnic groups in health outcomes, and these are influenced by both unequal social treatment and genetic differences to various degrees [38, 43]. The pattern of assortative mating found here suggests that these differences will persist for two reasons. First, the socially defined groups are being maintained and reinforced most strongly in the most diverse and culturally tolerant areas. Second, assortative mating will both preserve any allele frequency differences that exist between racial and ethnic groups that are associated with disease, and also bias individual genotypes towards homozygosity making disease phenotypes more likely to be expressed [36, 37].

More broadly, these findings suggest that gene-culture co-evolution is an on going process and that culture is affecting and structuring genomic allele frequencies on a massive scale [9, 10]. Indeed, cultural variation in the US explains the patterns of genetic diversity of more than 300 million people across a geographic area greater than 9.8 million square kilometres. Most research into human evolution has focused on understanding the past migrations and selective pressures that have shaped human genetic diversity [44]. The eminent geneticist Steve Jones has speculated that human evolution has stopped: “*World-wide, all populations are becoming connected and the opportunity for random change is dwindling. History is made in bed, but nowadays the beds are getting closer together. We are mixing into a global mass, and the future is brown*” [45]. This study provides strong evidence that this is not the case. Instead, cultural forces are structuring genetic diversity on a massive scale and assortative mating is maintaining diversity despite the great connectivity of peoples. Indeed, culture may be the predominant evolutionary force impacting human biology going forward.

## MATERIALS AND METHODS

### US Census demography

The US Census is performed every ten years and provides county level demographic data broken down by racial and ethnic membership for every US resident regardless of legal status [46]. In the most recent Census (2010) respondents were categorised as ‘Hispanic’, or ‘non-Hispanic’ ‘white’, ‘African American’, ‘Asian’, ‘Native American’, or ‘Pacific Islander’ alone or mixed, and numbers of each are reported for each county. The Census Bureau reports data for each year from 2010 to 2017, with 2010 being the raw data from the 2010 Census, and subsequent years being extrapolations from a model that considers births, deaths, and migration. I used data reported for 2016 in order to correspond with the 2016 US Presidential election. The demographic data were parsed into five study categories to best match up with the available genomic reference populations: European Americans, African Americans, Asian Americans, Native Americans, Pacific Islanders, and Hispanic Americans. These study categories were matched to the Census categories as follows: European Americans included ‘non-Hispanic white’, African Americans included ‘non-Hispanic African American’, Asian Americans included ‘non-Hispanic Asian’, Native Americans included ‘non-Hispanic Native American’, Pacific Islander included ‘non-Hispanic Pacific Islander’, and Hispanic included the sum of all ‘Hispanic’ categories.

### Genomic reference populations

A nationwide survey of current US genetic diversity found that genomic ancestry correlates highly with ethnicity as reported in the US Census [5]. As such, I used genetic reference populations to estimate genomic allele frequencies for each of the demographic categories included in the US Census. European American (CEU, N=99), African American (ASW, N=61), Hispanic American (MXL, N=52), and Asian American (CHB + GIH, N=206) references were taken from Phase 3 of the 1000 Genomes Project [4]. African Americans are known to vary regionally in their proportions of African and European admixture. African Americans from the US South generally have the highest amounts of African ancestry while those from the Southwest are known to have relatively more European ancestry [5]. The African American sample used here comes from the American Southwest and is therefore likely to have less African ancestry and therefore less genetic diversity than African Americans from elsewhere in the US. This is conservative relative to the hypotheses being tested here because it will tend to underestimate the genetic diversity of areas with high numbers of African Americans. Hispanics are a genetically diverse ethnicity, with those from Central America having more Native American ancestry and less European ancestry relative to those from the Caribbean [47]. The Mexican Americans in Los Angeles (MXL) sample was chosen alone to represent Hispanic American diversity because the greatest density of Hispanic Americans are located along the US-Mexico border (**Figure 1**). Limiting the Hispanic reference to the MXL is conservative with respect to the hypothesis being tested by potentially underestimating diversity in counties with Hispanic populations derived from the Caribbean. The Han Chinese (CHB) and Gujarati (GIH) were combined to represent the US Asian American population who broadly consist of people from South and East Asia. The Native American (Mayan + Pima + Karitiana, N=37) reference population was taken from the Human Genome Diversity Project (HGDP) samples genotyped with the Axiom Human Origins SNP chip [29]. Finally, the Pacific Islander reference population was a combination of the 1000 Genomes (JPT) and HGDP Japanese samples (N=132). The US Pacific Islander population are predominantly Native Hawaiians for which no genomic reference population is available. The Japanese samples were chosen as the best available substitute as an Island population who are relatively closely related to Polynesians [48]. Although not ideal, error from the use of Japanese as a reference for Pacific Islanders is unlikely to influence the findings, since Pacific Islanders only make up 0.17% of the US population (565,853 people in 2016) (**Figure 1**).

### Unbiased genomic SNP set

To reliably compare genetic diversity between structured groups it is imperative to use genetic markers for which the discovery was not biased towards any particular population [29]. The Axiom Human Origins SNPs from Panel 13 were discovered by sequencing a Denisovan and San individual [29], who are both expected to be equally distantly related to all of the reference study populations included in this study [49], and thus should have equal ascertainment. The complete 1000 Genomes Phase 3 data were downloaded and then filtered to only retain individuals from the reference populations and SNPs from the Axiom Human Origins Panel 13 using PLINK v1.9 [50]. The complete HGDP data were downloaded and filtered similarly. The phase of the HGDP data was then checked against the 1000 Genomes data using a custom Perl script, and SNPs out of phase were flipped using PLINK v1.9. The two datasets were then merged with PLINK v1.9 and individuals and SNPs with more than 5% missing data were excluded. The final dataset had 587 individuals and 46,154 SNPs with a genotyping rate of 0.971.

### Genetic diversity

Genetic diversity was estimated for each US county from county-level SNP allele frequencies which were in turn estimated from genomic reference population allele frequencies weighted by the county demography. Reference population allele frequencies were calculated using the PLINK –freq command. The county level frequency (*f_cnty_*) of each SNP was calculated as follows,

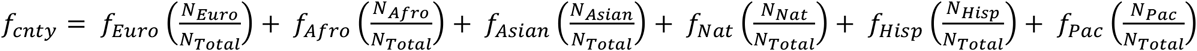

where *f_Euro_*, *f_Afro_*, *f_Asian_*, *f_Nat_*, *f_Hisp_*, *f_pac_* are the allele frequencies in the European American, African American, Asian American, Native American, Hispanic American, and Pacific Islander reference populations respectively, *N_Euro_*, *N_Afro_*, *N_Asian_*, *N_Nat_*, *N_Hisp_*, *N_pac_*, are the county population sizes of European Americans, African Americans, Asian Americans, Native Americans, Hispanic Americans, and Pacific Islanders respectively, and *N_Total_* is the total county population size.

Expected heterozygosity (*H_e_*) was then used as a measure of genetic diversity for each county [30]. Expected heterozygosity describes the probability that an individual from a population is heterozygous for any given locus. Heterozygosity (*h*) for each SNP for each county was calculated as follows.

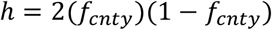

Genomic *H_e_* for each county was calculated from the individual SNP heterozygosities as follows.

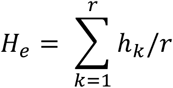

A custom Perl script was used to automate these calculations for every SNP for each US county.

### Relationship between genetic diversity and demography

Theoretical curves describing twoway demographic mixes of European Americans and each minority group were found empirically. To do so 100 hypothetical counties were constructed for each two-way demographic mix, with the proportion of European Americans ranging from 0 to 1 in 0.01 increments. Genetic diversity (*H_e_*) for each of the 100 hypothetical counties was then calculated as above. These points were then plotted and joined as a curve (**Figure 1**).

### Cultural and genetic niche separation

In order to measure the amount of cultural and genetic niche separation within states I created the statistics *Q_C_* and *Q_G_* respectively. These measures describe the amount of separation within states between counties that were won by Clinton or Trump relative to the total amount of cultural or genetic variation in the state. Both *Q_C_* and *Q_G_* are expressed in the number of standard deviations separating counties won by either candidate. *Q_C_* was computed as follows:

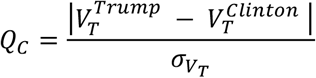

where 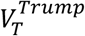 is the mean vote proportion for Donald Trump in counties that Trump won, 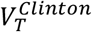 is the mean vote proportion for Donald Trump in counties that Clinton won, and *σ_V_T__* is the standard deviation in vote proportion for Donald Trump for all counties in the state. *Q_G_* was computed as follows:

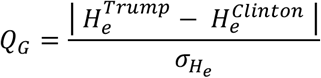

where 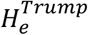 is the mean genetic diversity (*H_e_*) for counties won by Donald Trump, 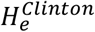 is the mean genetic diversity for counties won by Hillary Clinton, and *σ_H_e__* is the standard deviation in genetic diversity for all counties in the state.

### Assortative mating

The amount of assortative mating with respect to ethnicity was inferred from the US Census reports of whether people reported either a single or mixed ethnicity. This gives information about mate choice patterns from the parent generation of each individual in the census. An assortative mating index, *A*, was calculated to describe the relative departure from random mating with respect to ethnicity for each US county from the observed and expected frequencies of mixed ethnicity individuals in the census given the proportions of each census ethnic category. The expected frequency of mixed ethnicity individuals, *m_E_*, if mating were random was calculated as follows:

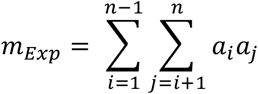

where *a_i_* is the frequency of the !th ethnic group in the population. The observed frequency of mixed individuals was calculated as follows:

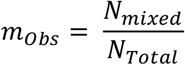

where *N_mixed_* is the number of people classed as having more than one ethnic affiliation in a county, and *N_Total_* is the total number of people in the county. ! was then calculated as follows:

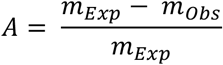

Values of *A* near zero indicate approximately random mating with respect to ethnicity. When *A* equals one, 100% of mate pairs are assortative with respect to ethnicity, and when *A* equals negative one, 100% of mate pairs are disassortative with respect to ethnicity. Values of greater than one or less than negative one are theoretically possible and would either reflect migration into or out of the county, or family structure within the data. These sources of bias are more likely to influence estimates of assortative mating in highly homogenous counties where the expected number of mixed individuals is very low.

### Statistical analyses and plots

Statistical analyses were performed as described in the main text using R [51]. Data were plotted using ggplot2 [52] for R. Maps were plotted using urbnmapr [53] and ggplot2 for R.

## ACKNOWLEDGMENTS

This research grew out of a talk for the Spring 2019 Symposium of the London Centre for Ecology and Evolution hosted by Roehampton University and I am thankful to Isabel Magalhaes for inviting me to participate in this symposium.

## REFERENCES

1. Leslie S, Winney B, Hellenthal G, Davison D, Boumertit A, Day T, et al. The fine-scale genetic structure of the British population. Nature. 2015;519(7543):309–14. doi: 10.1038/nature14230. PubMed PMID: 25788095.

2. Winney B, Boumertit A, Day T, Davison D, Echeta C, Evseeva I, et al. People of the British Isles: preliminary analysis of genotypes and surnames in a UK-control population. Eur J Hum Genet. 2012;20(2):203–10. doi: 10.1038/ejhg.2011.127. PubMed PMID: 21829225; PubMed Central PMCID: PMCPMC3260910.

3. Cann HM, de Toma C, Cazes L, Legrand MF, Morel V, Piouffre L, et al. A human genome diversity cell line panel. Science. 2002;296(5566):261–2. PubMed PMID: 11954565.

4. Durbin RM, Abecasis GR, Altshuler DL, Auton A, Brooks LD, Gibbs RA, et al. A map of human genome variation from population-scale sequencing. Nature. 2010;467(7319):1061–73. Epub 2010/10/29. doi: nature09534 [pii] 10.1038/nature09534. PubMed PMID: 20981092.

5. Bryc K, Durand EY, Macpherson JM, Reich D, Mountain JL. The Genetic Ancestry of African Americans, Latinos, and European Americans across the United States. American journal of human genetics. 2015;96(1):37–53. doi: 10.1016/j.ajhg.2014.11.010. PubMed PMID: 25529636.

6. Thiessen D, Gregg B. Human assortative mating and genetic equilibrium: An evolutionary perspective. Ethology and Sociobiology. 1980;1(2):111–40. doi: http://dx.doi.org/10.1016/0162-3095(80)90003-5.

7. Vandenberg SG. Assortative mating, or who marries whom? Behavior Genetics. 1972;2(2-3):127–57. doi: 10.1007/BF01065686.

8. Mesoudi A. Migration, acculturation, and the maintenance of between-group cultural variation. PLoS ONE. 2018;13(10):e0205573. doi: 10.1371/journal.pone.0205573. PubMed PMID: 30325943; PubMed Central PMCID: PMCPMC6191118.

9. Durham WH. Coevolution: Genes Culture and Human Diversity. Stanford, CA: Stanford University Press; 1991.

10. Laland KN, Odling-Smee J, Myles S. How culture shaped the human genome: bringing genetics and the human sciences together. Nature reviews. 2010;11(2):137–48. Epub 2010/01/20. doi: 10.1038/nrg2734. PubMed PMID: 20084086.

11. Laland KN, Odling-Smee J, Feldman MW. Cultural niche construction and human evolution. Journal of evolutionary biology. 2001;14(1):22–33. doi: 10.1046/j.1420-9101.2001.00262.x. PubMed PMID: 29280584.

12. Tishkoff SA, Reed FA, Ranciaro A, Voight BF, Babbitt CC, Silverman JS, et al. Convergent adaptation of human lactase persistence in Africa and Europe. Nat Genet. 2007;39(1):31–40. PubMed PMID: 17159977.

13. Perry GH, Dominy NJ, Claw KG, Lee AS, Fiegler H, Redon R, et al. Diet and the evolution of human amylase gene copy number variation. Nat Genet. 2007;39(10):1256–60. PubMed PMID: 17828263.

14. Ilardo MA, Moltke I, Korneliussen TS, Cheng J, Stern AJ, Racimo F, et al. Physiological and Genetic Adaptations to Diving in Sea Nomads. Cell. 2018;173(3):569–80 e15. doi: 10.1016/j.cell.2018.03.054. PubMed PMID: 29677510.

15. Liu H, Prugnolle F, Manica A, Balloux F. A geographically explicit genetic model of worldwide human-settlement history. American journal of human genetics. 2006;79(2):230–7. PubMed PMID: 16826514.

16. Manica A, Prugnolle F, Balloux F. Geography is a better determinant of human genetic differentiation than ethnicity. Human genetics. 2005;118(3-4):366–71. PubMed PMID: 16189711.

17. Prugnolle F, Manica A, Balloux F. Geography predicts neutral genetic diversity of human populations. Curr Biol. 2005;15(5):R159–60. PubMed PMID: 15753023.

18. Manica A, Amos W, Balloux F, Hanihara T. The effect of ancient population bottlenecks on human phenotypic variation. Nature. 2007;448(7151):346–8. PubMed PMID: 17637668.

19. Betti L, Manica A. Human variation in the shape of the birth canal is significant and geographically structured. Proceedings. 2018;285(1889). doi: 10.1098/rspb.2018.1807. PubMed PMID: 30355714; PubMed Central PMCID: PMCPMC6234894.

20. Betti L, Balloux F, Amos W, Hanihara T, Manica A. Distance from Africa, not climate, explains within-population phenotypic diversity in humans. Proceedings. 2009;276(1658):809–14. doi: 10.1098/rspb.2008.1563. PubMed PMID: 19129123; PubMed Central PMCID: PMCPMC2664379.

21. Kirby KR, Gray RD, Greenhill SJ, Jordan FM, Gomes-Ng S, Bibiko HJ, et al. D-PLACE: A Global Database of Cultural, Linguistic and Environmental Diversity. PLoS ONE. 2016;11(7):e0158391. doi: 10.1371/journal.pone.0158391. PubMed PMID: 27391016; PubMed Central PMCID: PMCPMC4938595.

22. Erikson RS, McIver JP, Wright GC. State Political Culture and Public Opinion. American Political Science Review. 2014;81(3):797–813. Epub 08/01. doi: 10.2307/1962677.

23. Swauger J. Regionalism in the 1976 Presidential Election. Geographical Review. 1980;70(2):157–66. doi: 10.2307/214437.

24. Moore LM, Ovadia S. Accounting for Spatial Variation in Tolerance: The Effects of Education and Religion. Social Forces. 2006;84(4):2205–22. doi: 10.1353/sof.2006.0101.

25. Pilkington E, Jacobs B. Donald Trump: ban all Muslims entering the US. The Guardian. 2015.

26. Bobo LD. Racism in Trump’s America: reflections on culture, sociology, and the 2016 US presidential election. The British Journal of Sociology. 2017;68(S1):S85–S104. doi: 10.1111/1468-4446.12324.

27. Huang J, Jacoby S, Strickkland M, Lai KKR. Election 2016: Exit Polls. The New York Times. 2016.

28. Hersh ED, Nall C. The Primacy of Race in the Geography of Income-Based Voting: New Evidence from Public Voting Records. American Journal of Political Science. 2016;60(2):289–303. doi: 10.1111/ajps.12179.

29. Patterson N, Moorjani P, Luo Y, Mallick S, Rohland N, Zhan Y, et al. Ancient admixture in human history. Genetics. 2012;192(3):1065–93. Epub 2012/09/11. doi: 10.1534/genetics.112.145037. PubMed PMID: 22960212; PubMed Central PMCID: PMC3522152.

30. Nei M. Estimation of average heterozygosity and genetic distance from a small number of individuals. Genetics. 1978;89:583–90.

31. MIT Election Data + Science Lab Cambridge, MA, USA.: Massachusets Institute of Technology; 2019 [cited 2019 07-2019]. Available from: https://electionlab.mit.edu/data.

32. Emerson FV. Geographic influences in American slavery. Bulletin of the American Geographical Society of New York (1901-1915). 1911;43(1):13. PubMed PMID: 125736196.

33. Mondak JJ, Canache D. Personality and Political Culture in the American States. Political Research Quarterly. 2013;67(1):26–41. doi: 10.1177/1065912913495112.

34. Trump’s tweet: What did he say and why’s he being criticised? The BBC. 2019 18/07/2019.

35. Burnham KP, Anderson DR. Multimodel inference - understanding AIC and BIC in model selection. Sociol Method Res. 2004;33(2):261–304. doi: 10.1177/0049124104268644. PubMed PMID: WOS:000224706300004.

36. Crow JF, Kimura M. An Introduction to Population Genetics Theory. New York: Harper and Row; 1970.

37. Crow JF, Felsenstein J. The effect of assortative mating on the genetic composition of a population. Eugen Q. 1968;15(2):85–97. PubMed PMID: 5702332.

38. Gravlee CC. How race becomes biology: embodiment of social inequality. American journal of physical anthropology. 2009;139(1):47–57. doi: 10.1002/ajpa.20983. PubMed PMID: 19226645.

39. Jorde LB, Wooding SP. Genetic variation, classification and ‘race’. Nat Genet. 2004;36(11 Suppl):S28–33. Epub 2004/10/28. doi: 10.1038/ng1435. PubMed PMID: 15508000.

40. Fremon DK. The Jim Crow laws and racism in American history. Berkeley Heights, NJ, USA.: Enslow Publishers; 2000.

41. Acharya A, Blackwell M, Sen M. The Political Legacy of American Slavery. The Journal of Politics. 2016;78(3):621–41. doi: 10.1086/686631.

42. Anderson E, Austin DW, Holloway CL, Kulkarni VA. The Legacy of Racial Caste: An Exploratory Ethnography. The Annals of the American Academy of Political and Social Science. 2012;642(1):25–42. doi: 10.1177/0002716212437337.

43. Bamshad M, Wooding S, Salisbury BA, Stephens JC. Deconstructing the relationship between genetics and race. Nature reviews. 2004;5(8):598–609. doi: 10.1038/nrg1401. PubMed PMID: 15266342.

44. Novembre J, Ramachandran S. Perspectives on human population structure at the cusp of the sequencing era. Annual review of genomics and human genetics. 2011;12:245–74. Epub 2011/08/02. doi: 10.1146/annurev-genom-090810-183123. PubMed PMID: 21801023.

45. Belluz J. Leading geneticist Steve Jones says human evolution is over. The Times. 2008 October 7, 2008.

46. US_Census_Bureau. County Population Totals and Components of Change: 2010–2018 http://www.census.gov: US Census Bureau; 2010. Available from: https://www.census.gov/data/tables/time-series/demo/popest/2010s-counties-total.html.

47. Bryc K, Velez C, Karafet T, Moreno-Estrada A, Reynolds A, Auton A, et al. Colloquium paper: genome-wide patterns of population structure and admixture among Hispanic/Latino populations. Proceedings of the National Academy of Sciences of the United States of America. 2010;107 Suppl 2:8954–61. Epub 2010/05/07. doi: 0914618107 [pii] 10.1073/pnas.0914618107. PubMed PMID: 20445096; PubMed Central PMCID: PMC3024022.

48. Friedlaender JS, Friedlaender FR, Reed FA, Kidd KK, Kidd JR, Chambers GK, et al. The Genetic Structure of Pacific Islanders. PLoS genetics. 2008;4(1):e19.

49. Meyer M, Kircher M, Gansauge M-T, Li H, Racimo F, Mallick S, et al. A High-Coverage Genome Sequence from an Archaic Denisovan Individual. Science. 2012.

50. Purcell S, Neale B, Todd-Brown K, Thomas L, Ferreira MA, Bender D, et al. PLINK: a tool set for whole-genome association and population-based linkage analyses. American journal of human genetics. 2007;81(3):559–75. Epub 2007/08/19. doi: S0002-9297(07)61352-4 [pii] 10.1086/519795. PubMed PMID: 17701901; PubMed Central PMCID: PMC1950838.

51. R_Development_Core_Team. R: A language and environment for statistical computing. Vienna, Austria: R Foundation for Statistical Computing; 2011.

52. Wickham H. ggplot2: elegant graphics for data analysis. New York, NY, USA: Springer; 2009.

53. Williams A, Ueyama K, Strochak S. Urbnmapr: How to create state and county maps easily in R. Urban Institute; 2018.

