## Supplementary Information for "Cultural variation in voting patterns reflects the landscape of US genetic diversity: A test of the cultural niche hypothesis"

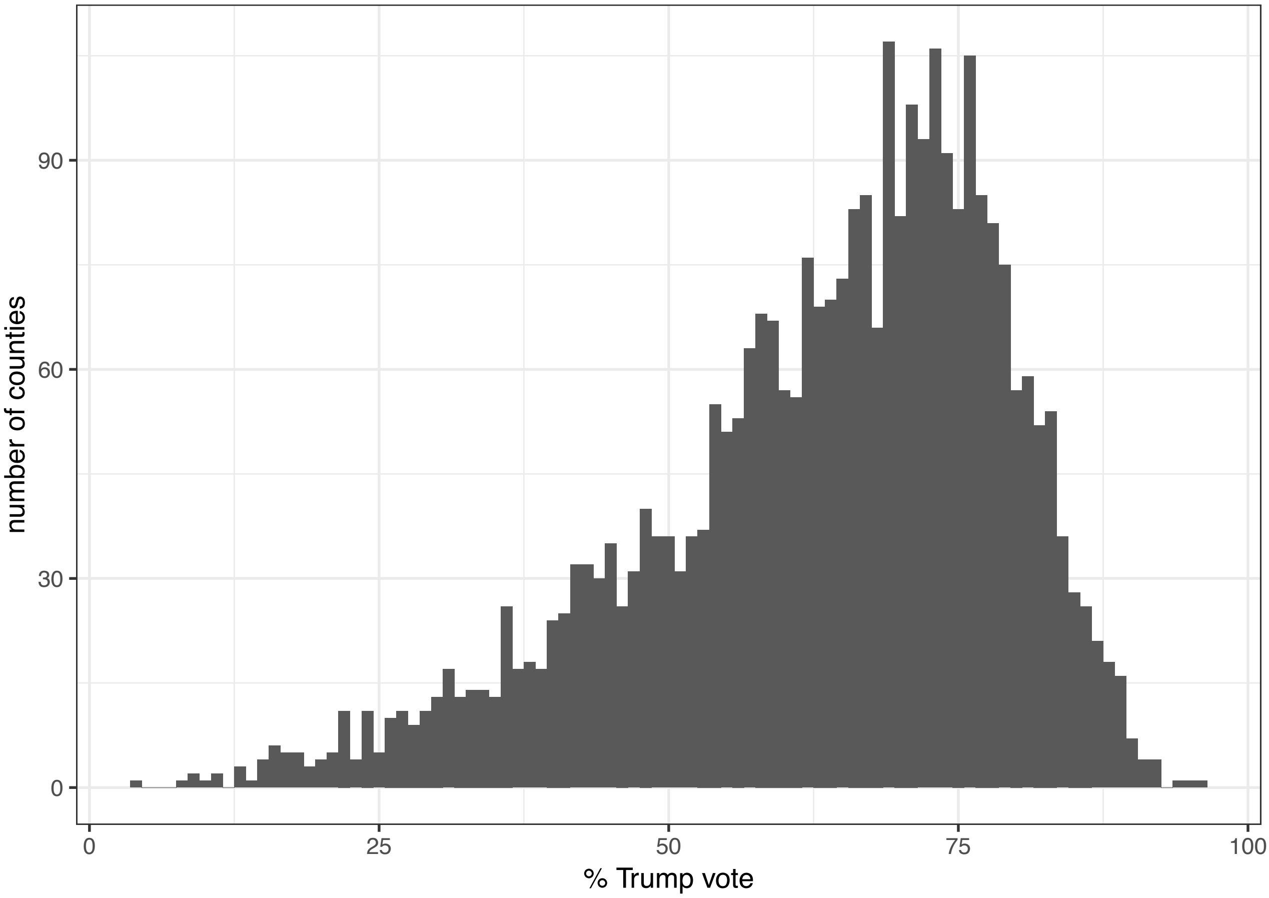


**Figure S1.** Histogram of county-level variation in 2016 US Presidential Election voting (excluding Alaska). The distribution is heavily skewed reflecting the 2,623 to 488 advantage of counties won by Donald Trump relative to Hillary Clinton.

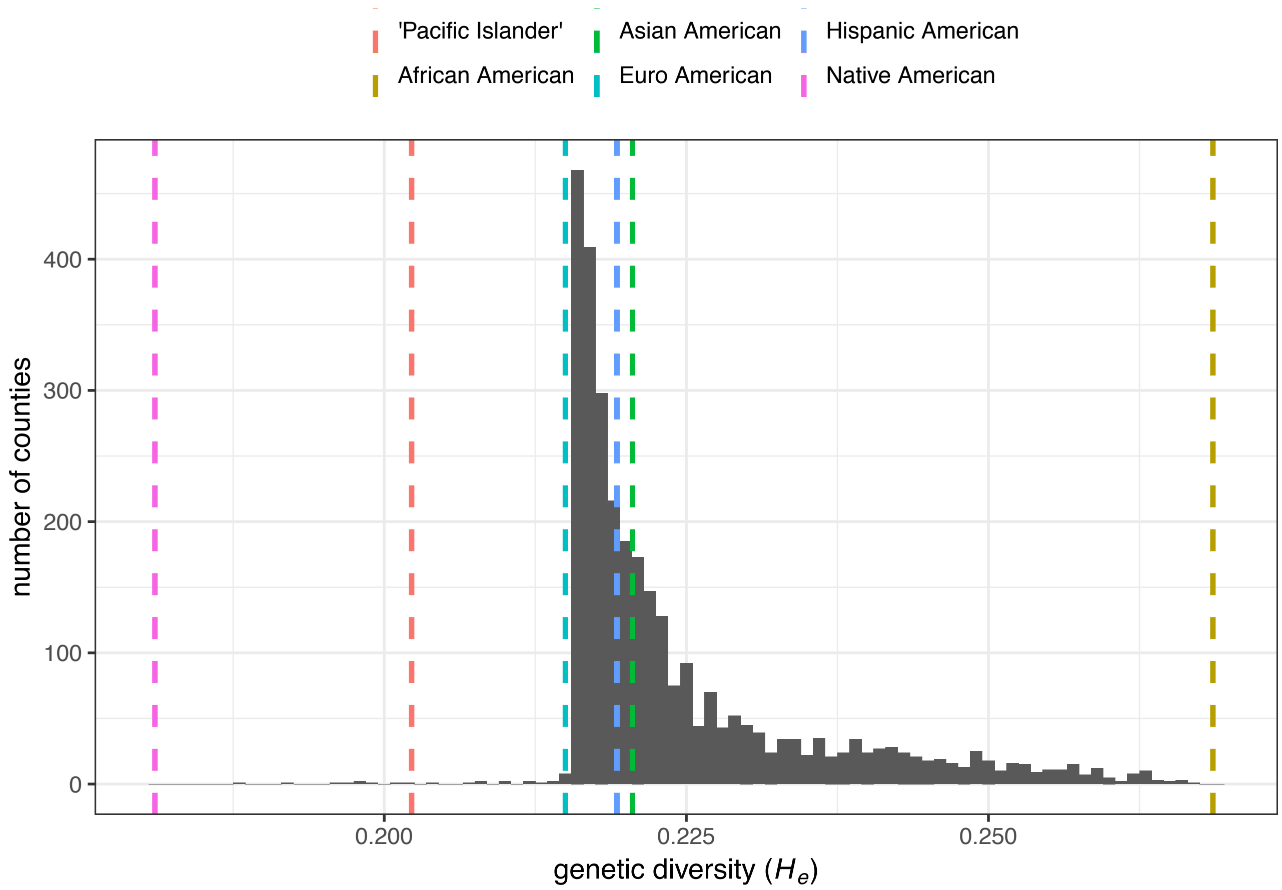


**Figure S2.** Histogram of US county-level genetic diversity. The vertical dashed lines repesent the genetic diversity of each demographic included in the 2010 US Census. Most counties have genetic diversity close to that of the majority demographic European Americans. Counties in the long right tail tend to have a high proportion of African Americans who are the most geneticaly diverse demographic. The small number of counties in the left tail have a high proportion of Native Americans who are the least genetically diverse demographic.

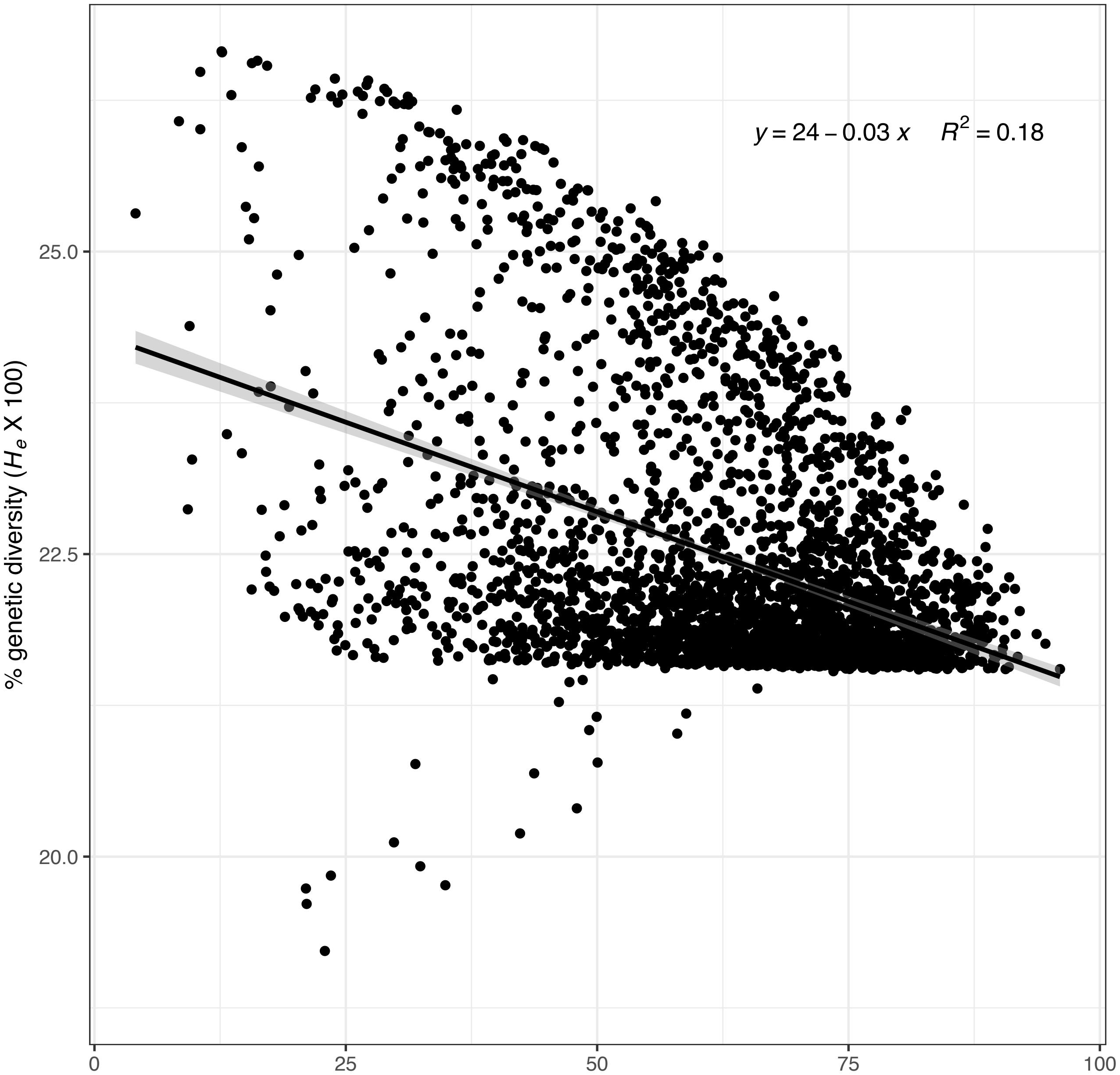


**Figure S3.** The relationship between county-level preference for Trump and genetic diversity, $H_e$ (excluding Alaska). A linear model found that nationwide, the 2016 US Presidential Election explains 18% of the variation in genetic diversity, with a 10% increase in voting for Donald Trump associated with a 0.3% decrease in genetic diversity ($F_{1,3109}$=689.1, $R^2$=0.18, p<2.2e-16, AIC=8751.5).

**Region: Northeast**

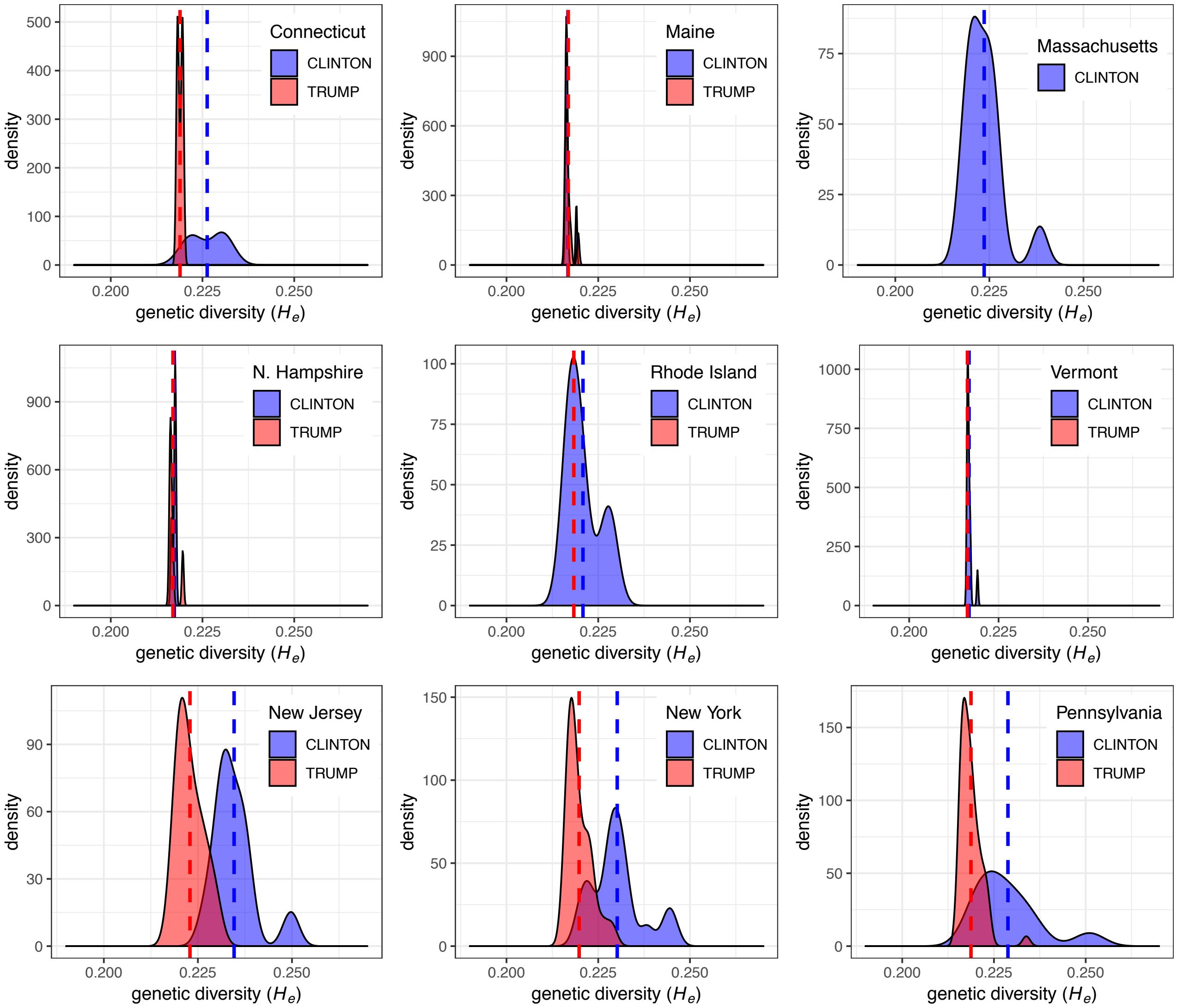


**Figure S4 (Northeast).** Density plots of county-level variation in genetic diversity by whether the county preferred Trump or Clinton for each US State (excluding Alaska). In the majority of US states counties that voted for Hillary Clinton have higher genetic diversity than counties that voted for Donald Trump.

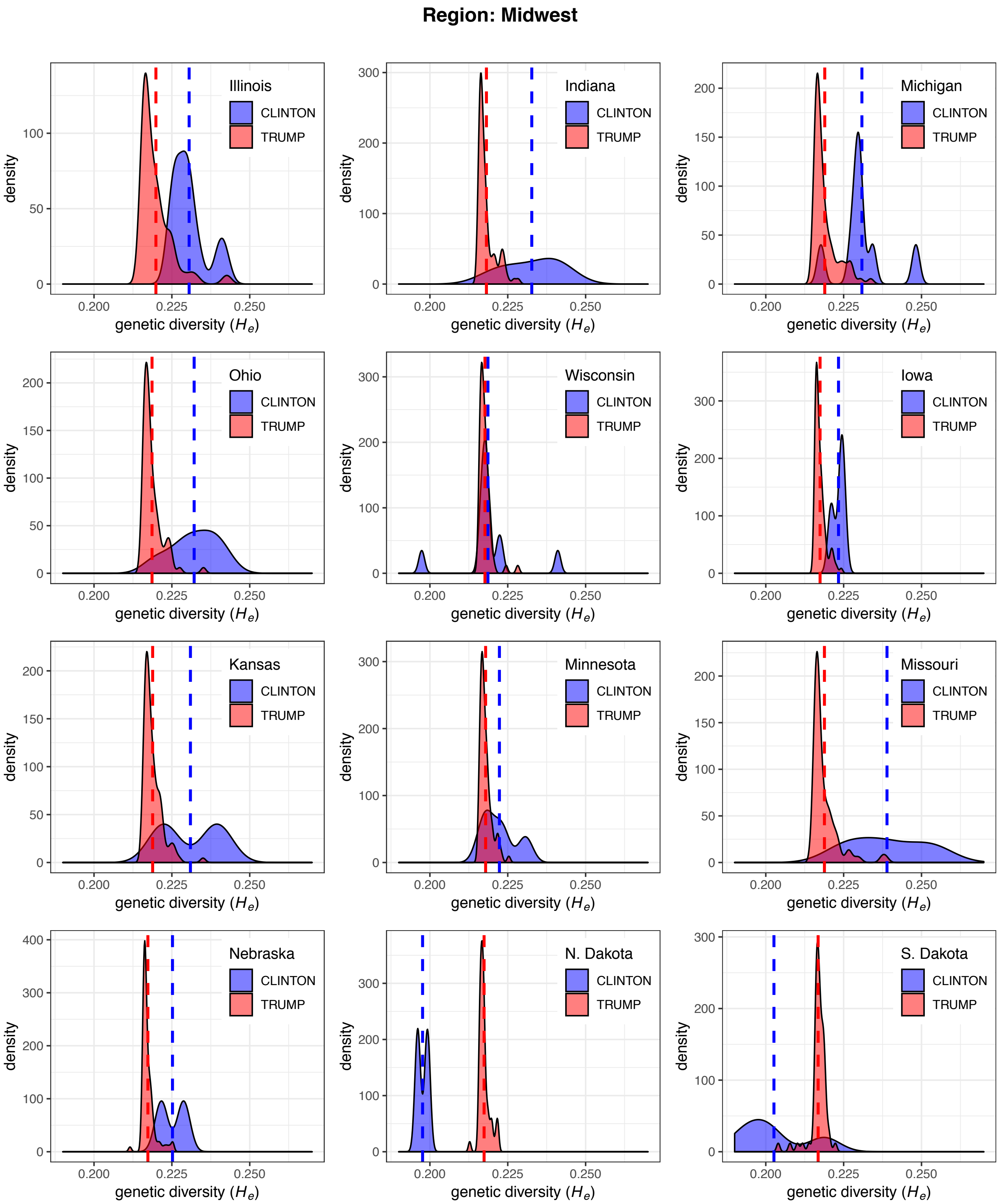


**Figure S4 (Midwest).** Density plots of county-level variation in genetic diversity by whether the county preferred Trump or Clinton for each US State (excluding Alaska). In the majority of US states counties that voted for Hillary Clinton have higher genetic diversity than counties that voted for Donald Trump.

**Region: South**

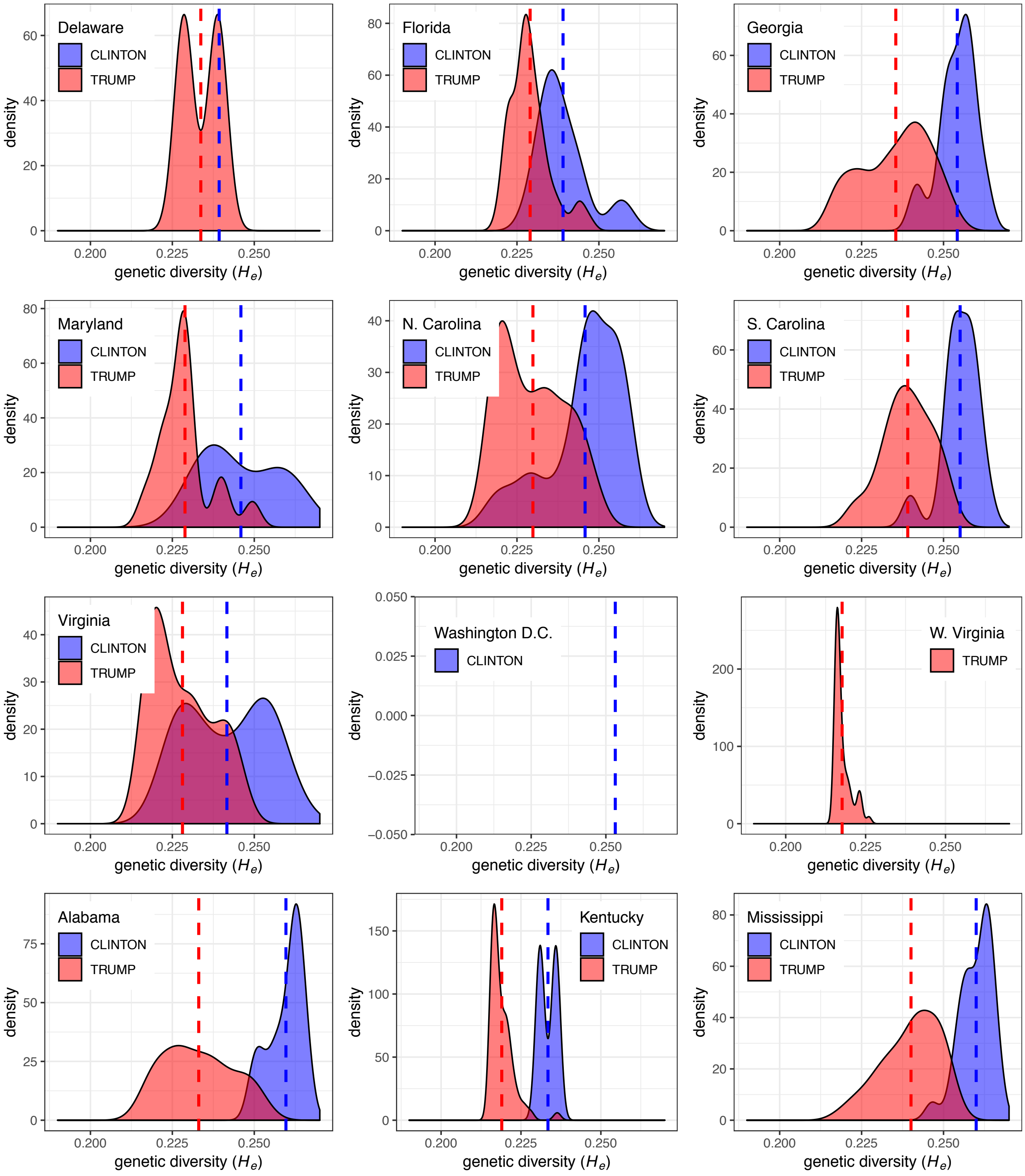


**Figure S4 (South).** Density plots of county-level variation in genetic diversity by whether the county preferred Trump or Clinton for each US State (excluding Alaska). In the majority of US states counties that voted for Hillary Clinton have higher genetic diversity than counties that voted for Donald Trump.

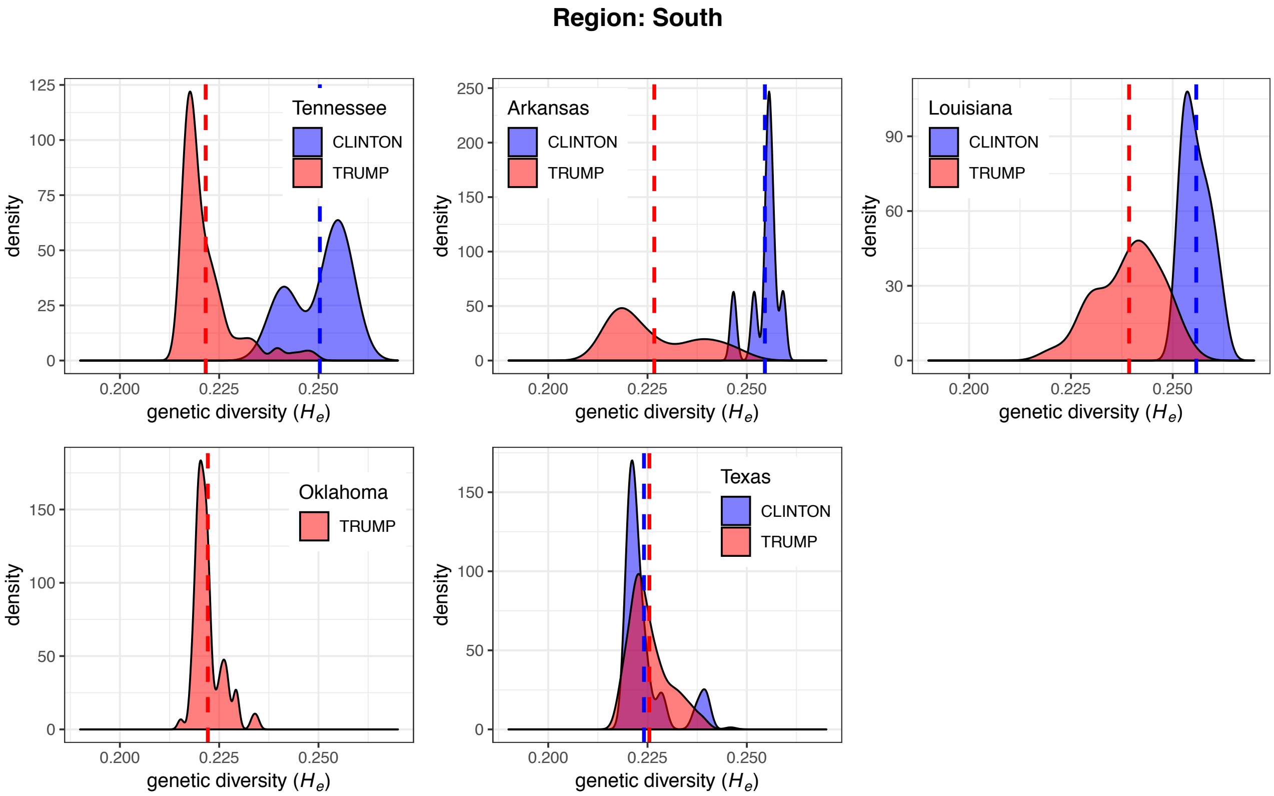


**Figure S4 (South continued).** Density plots of county-level variation in genetic diversity by whether the county preferred Trump or Clinton for each US State (excluding Alaska). In the majority of US states counties that voted for Hillary Clinton have higher genetic diversity than counties that voted for Donald Trump.

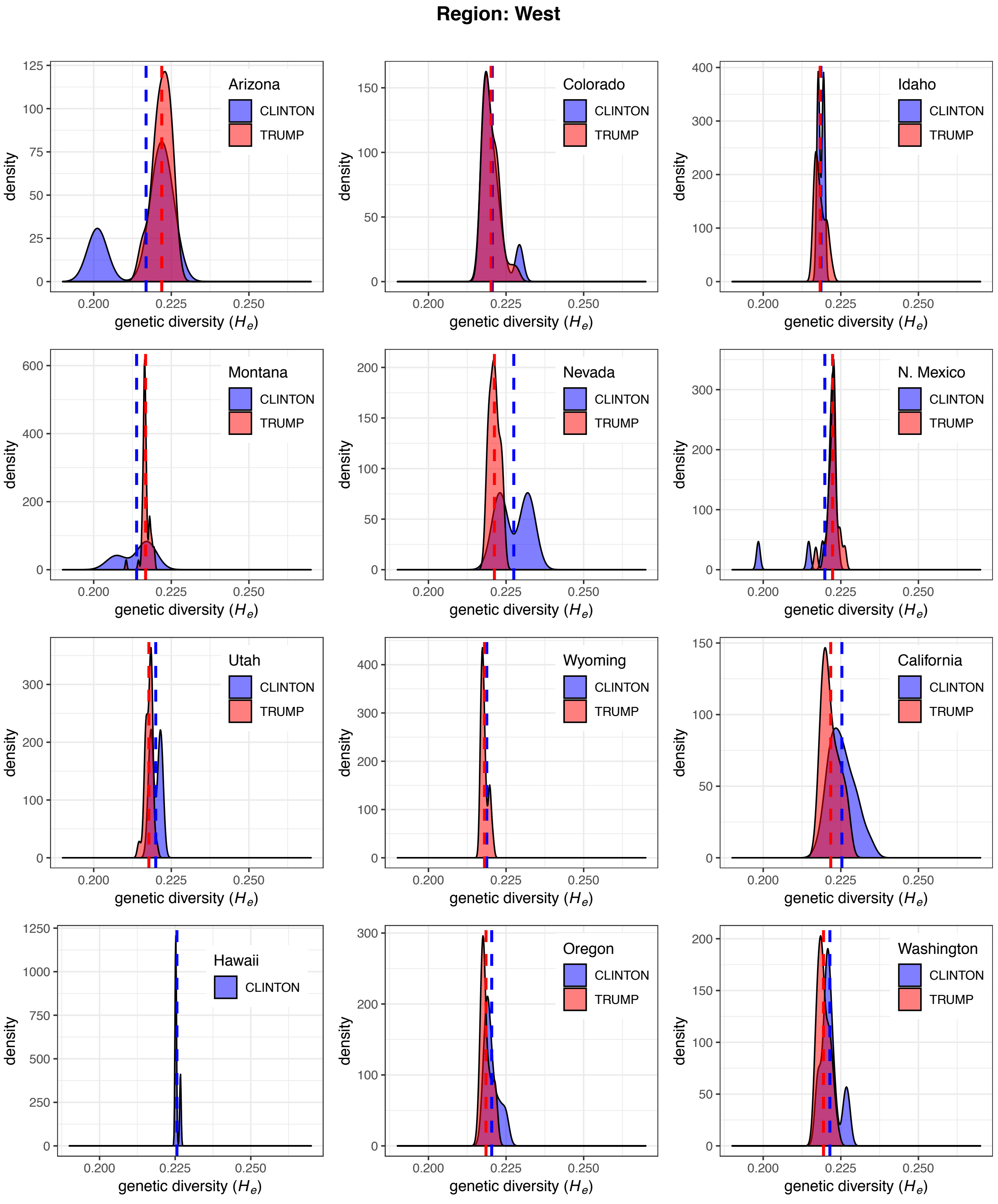


**Figure S4 (West).** Density plots of county-level variation in genetic diversity by whether the county preferred Trump or Clinton for each US State (excluding Alaska). In the majority of US states counties that voted for Hillary Clinton have higher genetic diversity than counties that voted for Donald Trump.

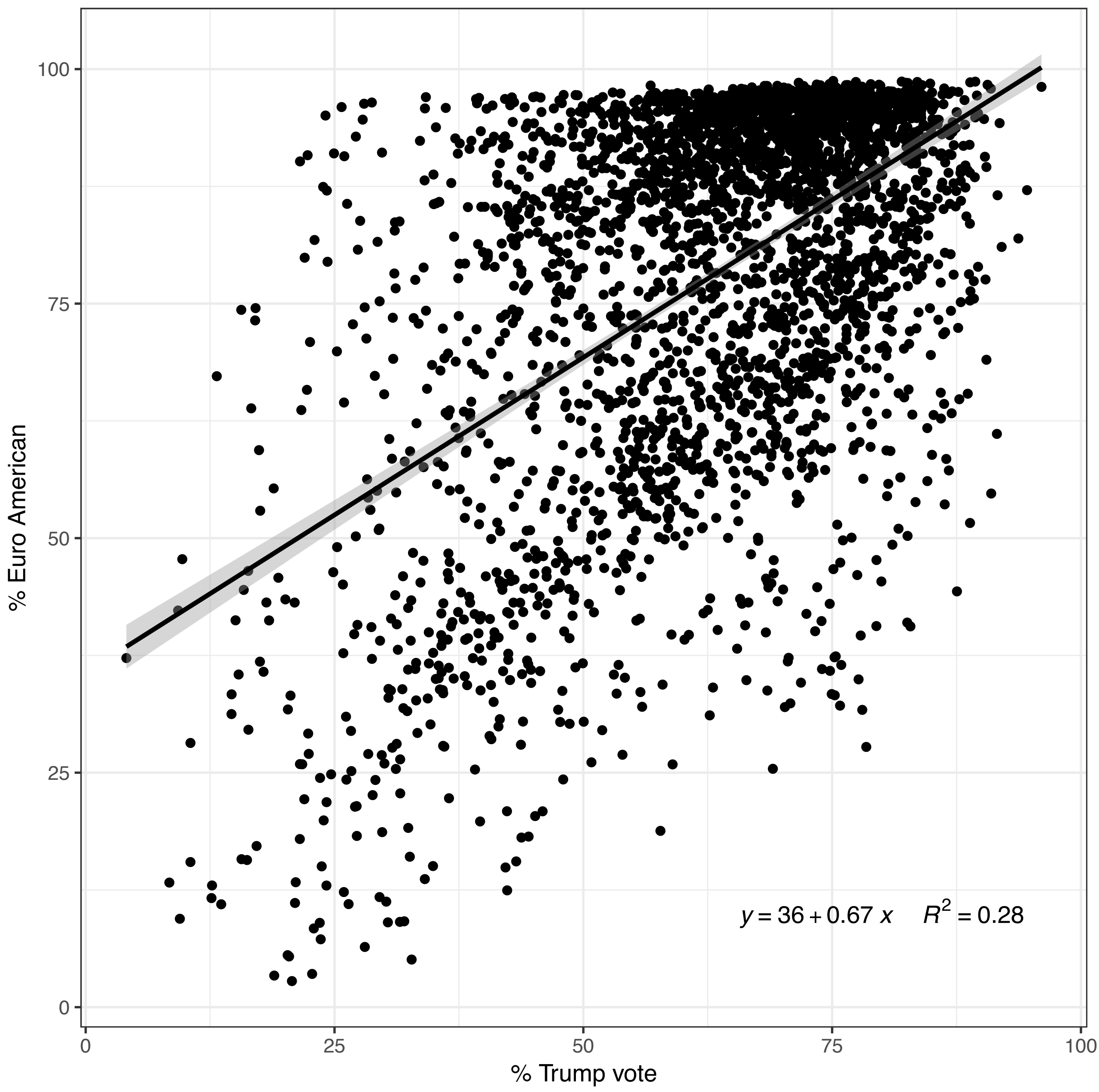


**Figure S5.** The relationship between US county-level demography and preference for Donald Trump (excluding Alaska). Nationwide variation in the 2016 US Presidential Election explains 28% of the variation in the percentage of European Americans in each county ($F_{1,3109}$=1192, $R^2$=0.28, p<2.2e-16, AIC=26,468.1). Every 10% increase in votes for Trump is associated with a 6.7% increase in the proportion of European Americans.

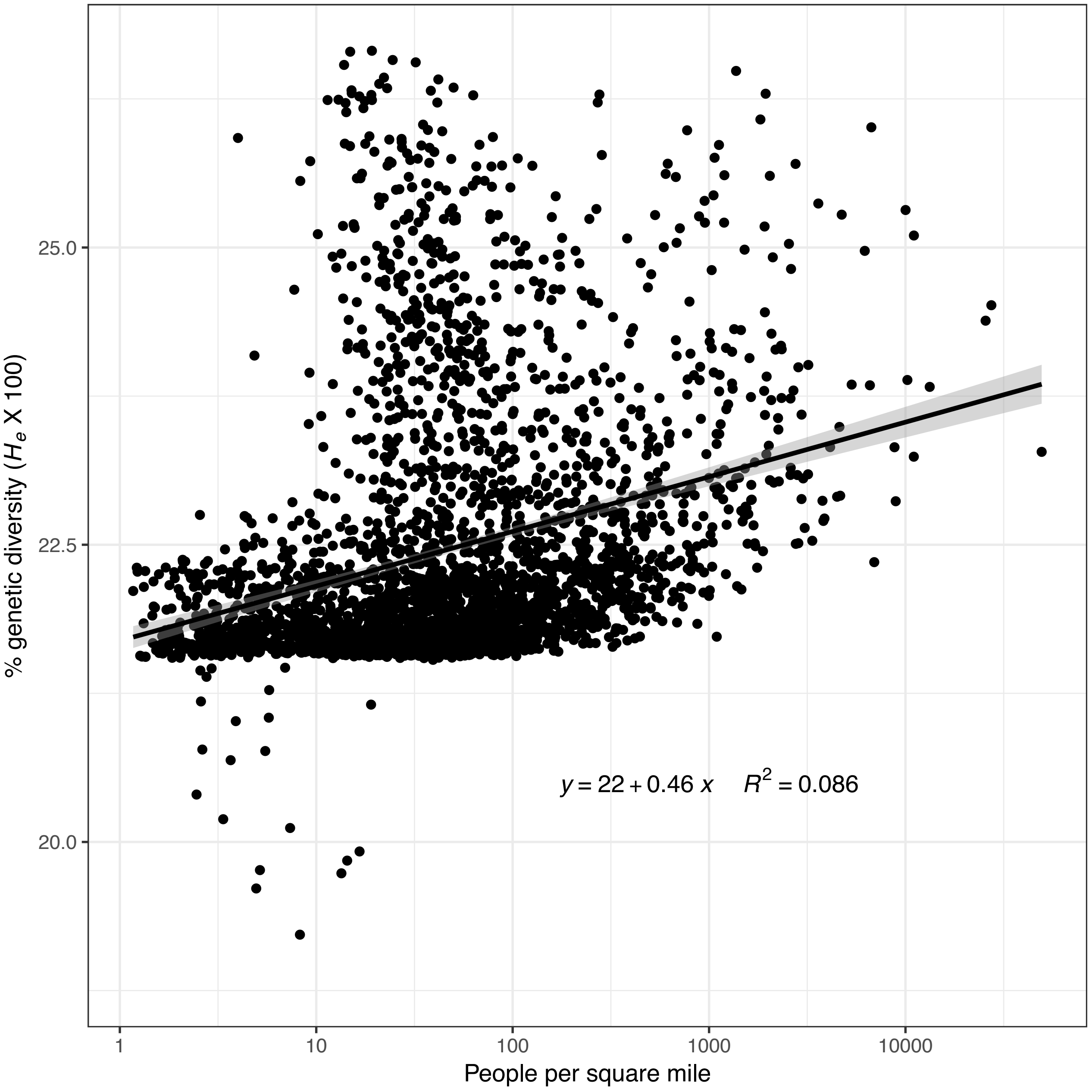


**Figure S6.** The relationship between genetic diversity and population density. Linear regression was used to model genetic diversity as a function of the $\log_{10}$ of population density ($F_{1,3109}$=292.2, $R^2$=0.086, p<2.2e-16, AIC=9095.0). Population density explains 9% of the variation in genetic diversity.

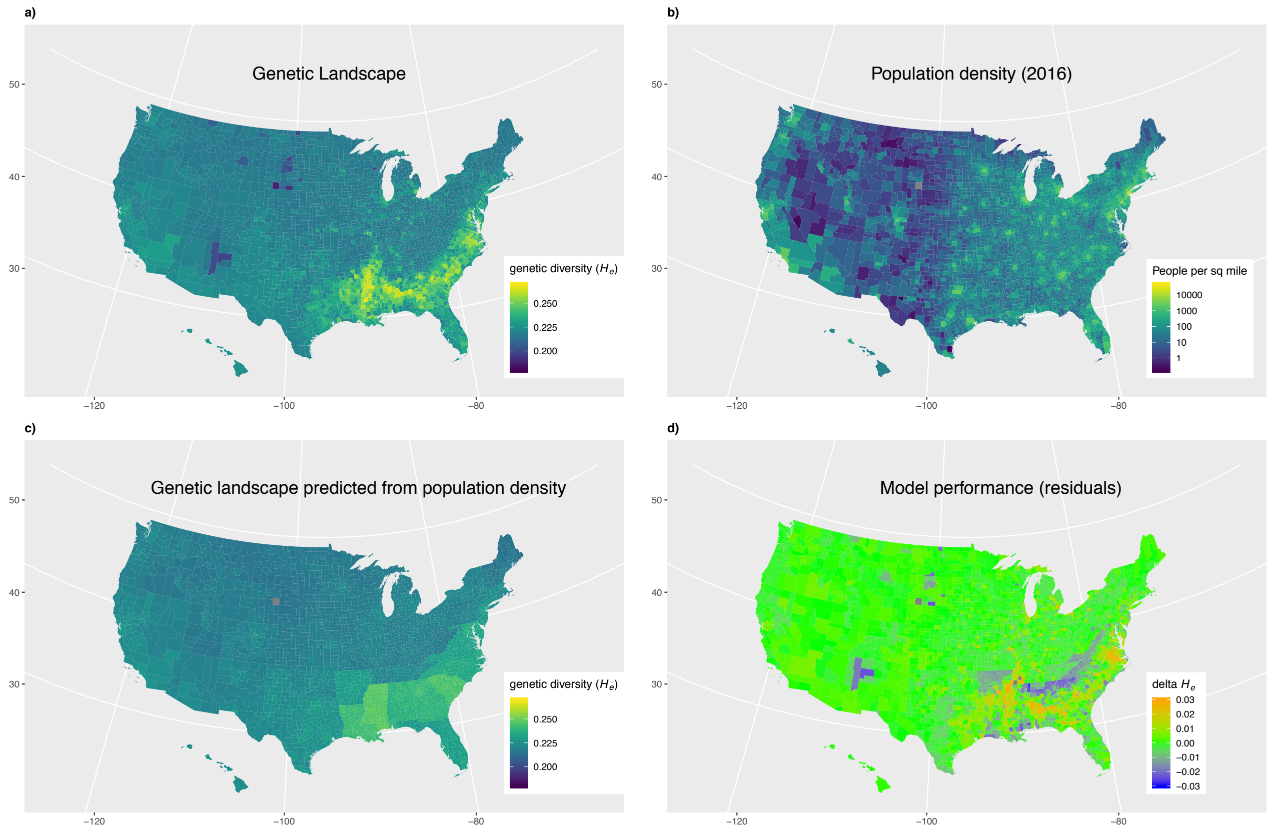


**Figure S7.** The relationship between population density and genetic diversity. The data are provided by the US Census Bureau for the year 2016. **a)** The landscape of genetic diversity ($H_e$). **b)** US county-level population density in people per square mile, **c)** The landscape of genetic diversity predicted from the $\log_{10}$ of population density controlling for state level variation. This model explains 57% of the variation in genetic diversity ($F_{50,3060}$=84.9, $R^2$=0.574, p<2.2e-16, AIC=6766.1), but only 2% more than a model that considers state level variation alone. **d)** The difference between observed and predicted genetic diversity. The model performs poorly in counties with very high and very low genetic diversity.

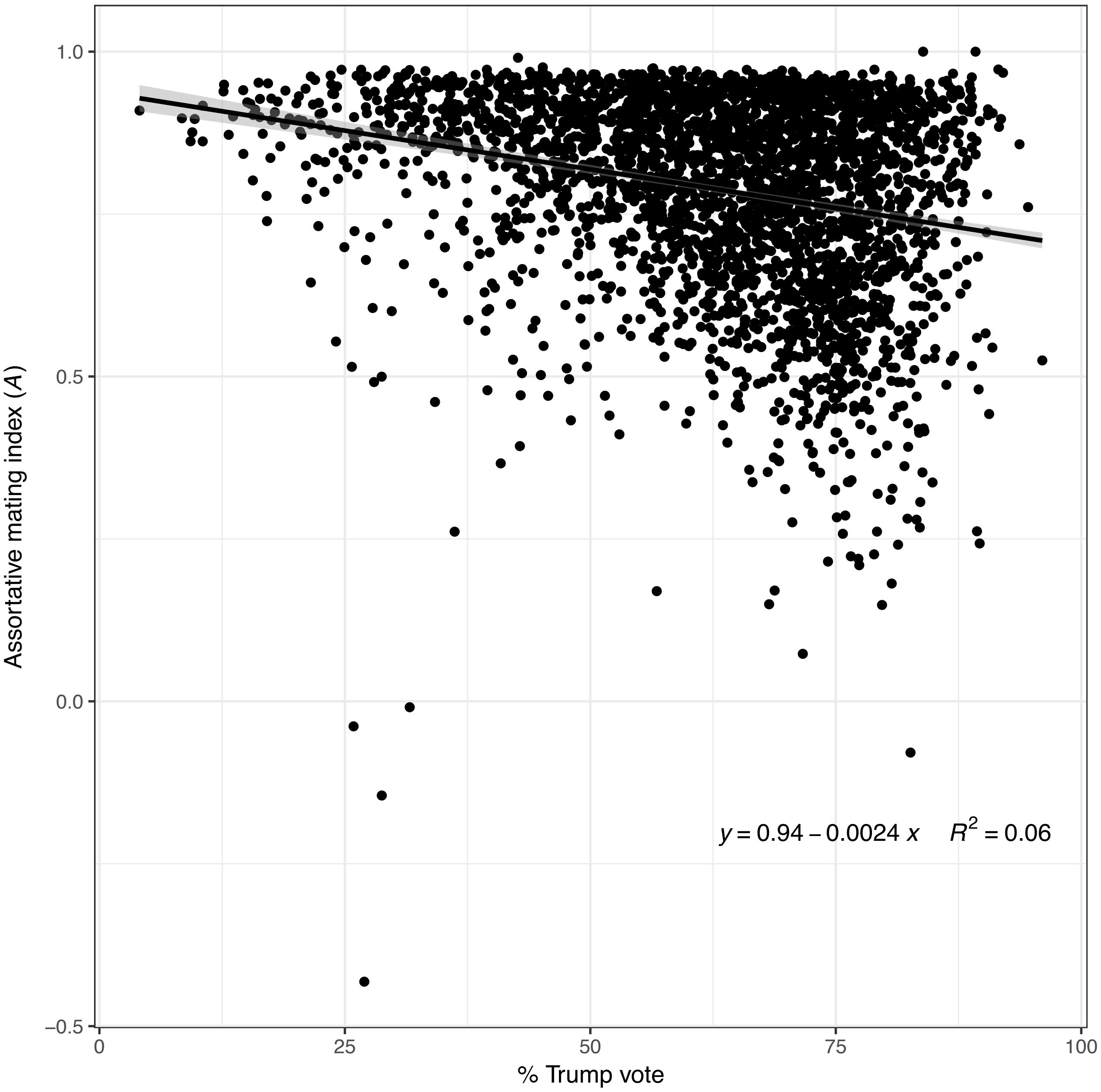


**Figure S8.** The relationship between assortative mating and preference for Donald Trump. There is a weak negative relationship between assortative mating and preference for Trump ($F_{1,3109}$=198.8, $R^2$=0.06, p<2.2e-16).
